# Bacterial Peptidoglycan Fragments Differentially Regulate Innate Immune Signaling

**DOI:** 10.1101/2020.09.03.278705

**Authors:** Klare L. Bersch, Kristen E. DeMeester, Rachid Zagani, Kimberly A. Wodzanowski, Hans-Christian Reinecker, Catherine L. Grimes

## Abstract

The human innate immune system responds to both pathogen and commensal bacteria at the molecular level using bacterial peptidoglycan (PG) recognition elements. Traditionally, synthetic and commercially accessible PG monosaccharide units known as muramyl dipeptide (MDP) and *N*-glycolyl MDP (ng-MDP) have been used to probe the mechanism of innate immune activation of pattern recognition receptors (PRRs) such as NOD-like receptors (NLRs). However, bacterial PG is a dynamic and complex structure, with various chemical modifications and trimming mechanisms that result in the production of disaccharide containing elements. These molecules pose as attractive targets for immunostimulatory screening; however, studies are limited due to their synthetic accessibility. Inspired by disaccharide containing compounds produced from the gut microbe, *Lactobacillus acidophilus*, a robust and scalable chemical synthesis of PG-based disaccharide ligands was implemented. Together with a monosaccharide PG library, compounds were screened for their ability to stimulate proinflammatory genes in bone marrow derived macrophages (BMDMs). The data reveal a diverse gene induction pattern between monosaccharide and disaccharide PG units, suggesting that PG innate immune signaling is more complex than a one-activator-one pathway program, as biologically relevant fragments induce distinct transcriptional programs. These disaccharide molecules will serve as critical immunostimulatory tools to more precisely define specialized innate immune regulatory mechanisms that distinguish between commensal and pathogenic bacteria residing in the microbiome.

## Introduction

The human body is responsible for maintaining ∼39 trillion bacterial cells^1-4^ that constitute the microbiome^5-6^. The gut microflora is one area of the body teeming with hundreds of species of bacterial cells^7-9^. The bacteria in the gastrointestinal (GI) tract are benefactors to the human host, performing essential biological chemical transformations^10, 11-13^ and producing key essential vitamins and amino acids^14^. While many of these organisms serve to maintain a healthy state for the human host, bacterial pathogenesis disrupts this symbiotic relationship. Dysbiosis in the human microbiome can lead to a variety of inflammatory diseases, including ulcerative colitis and Crohn’s disease (CD), rheumatoid arthritis, GI cancer, and asthma^15^. Therefore, the host-microbiome interface poses as an attractive target for therapeutic intervention^16^. In order to develop novel immunotherapies and antibiotics, it is critical to fully understand the molecular mechanisms by which Nature recognizes and responds to bacteria.

Humans have developed host-defense mechanisms to combat infectious diseases including the innate immune system, the body’s first line of defense against invading pathogens such as bacteria^17^. Pattern recognition receptors (PRRs)^18-22^ are programmed in this system to interact with essential components of bacterial cells such as flagella, lipopolysaccharide (LPS), lipoteichoic acids, and bacterial cell wall peptidoglycan (PG) components (Scheme 1)^1, 23-24^. Since the pioneering PRR discovery, multiple families of PRRs have been classified such as: Toll-like receptors (TLRs), C-type lectin receptors (CLRs), NOD-like receptors (NLRs), RIG-I-like receptors (RLRs), AIM2-like receptors (ALRs), Peptidoglycan binding proteins (PGBPs), SLAM family (SLAMF) and OAS-like receptors (OLRs)^15-17^. The broad specificity of PRRs is fascinating; how this system provides a tuned innate immune response towards pathogens and distinguishes symbiotic microorganisms that constitute the microbiome^25^ is not fully understood.

The bacterial microbe associate molecular pattern (MAMP) peptidoglycan (PG) is sensed by a variety of PRRs including NOD1 and NOD2, NLRP3, NLRP1 and PGBPs (Scheme 1)^26-32^. These ligands are small fragments derived from a large PG polymer that surrounds the bacterial cell, consisting of repetitive units of β-1,4 linked *N-*acetylglucosamine (GlcNAc) and *N-*acetylmuramic acid (MurNAc) with short peptide chains containing both _L_ and _D_ amino acids present on the muramic acid residue (Figure 1). Although the bacterial cell wall is highly conserved among bacterial species, differences arise (Figure 1)^33-34^. Variations in cross-linking (3-3 vs. 3-4 vs. 2-4 linkages), as well as substitution of amino acids, primarily at the 3^rd^ position (such as meso-diaminopimelic acid (m-DAP), _L_ -Lys, _L_ -Orn, _L_ -Ala, _L_ -Glu, _L_ -Homoserine), are observed in both Gram-positive and Gram-negative bacteria (Figure 1, shown in blue) ^33^. Modifications to the carbohydrate backbone of PG have also been identified in a variety of bacterial species including: *N*-deacetylation in *Listeria*^*35*^, *O-*acetylation in *Helicobacter pylori*^*36*^, *N-*glycolylation in *Mycobacterium tuberculousis*^*37*^, Muramic δ-lactam in *Bacillus subtilis*^*38*^, and 1,6-Anhydro^39^ limited all block PG lytic enzymatic digestion, leaving 1,6-Anhydro as the lytic product (Figure 1, shown in red). From this knowledge of PG complexity, one can easily imagine a pool of tunable immunostimulatory fragments critical for mediating host-pathogen interactions. However, the small molecule details in this signaling landscape is incomplete due to a limited amount of biologically significant MAMP-PG chemical probes.

**Scheme 1.**
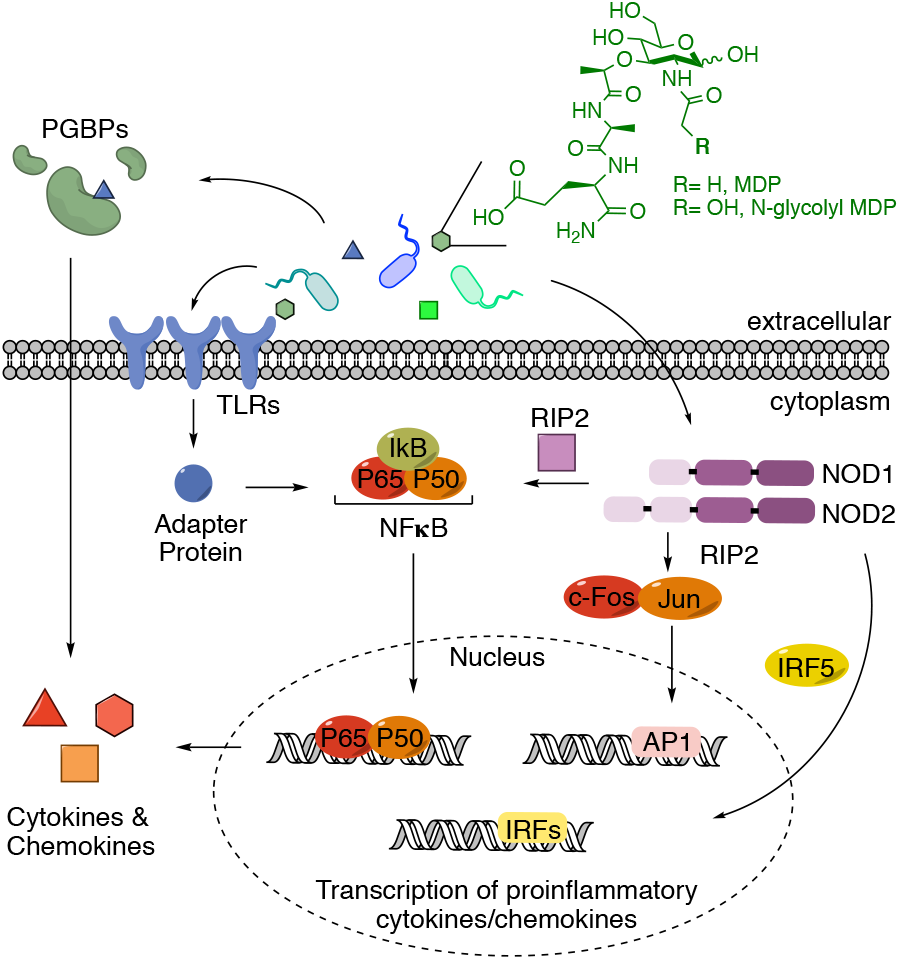
PRR signaling. Common activation pathways of proinflammatory cytokine and chemokine production upon stimulation by molecular signatures known as pathogen associated molecular patterns (PAMPS)^1-2^. In the advent of the microbiome, due to similarities between pathogenic and non-pathogenic organisms, these signals are now referred to as microbe associate molecular patterns (MAMPS)^3^. Here the model of activation by PG (and synthetic mimics, MDP and ng-MDP) is shown.

**Figure 1.**
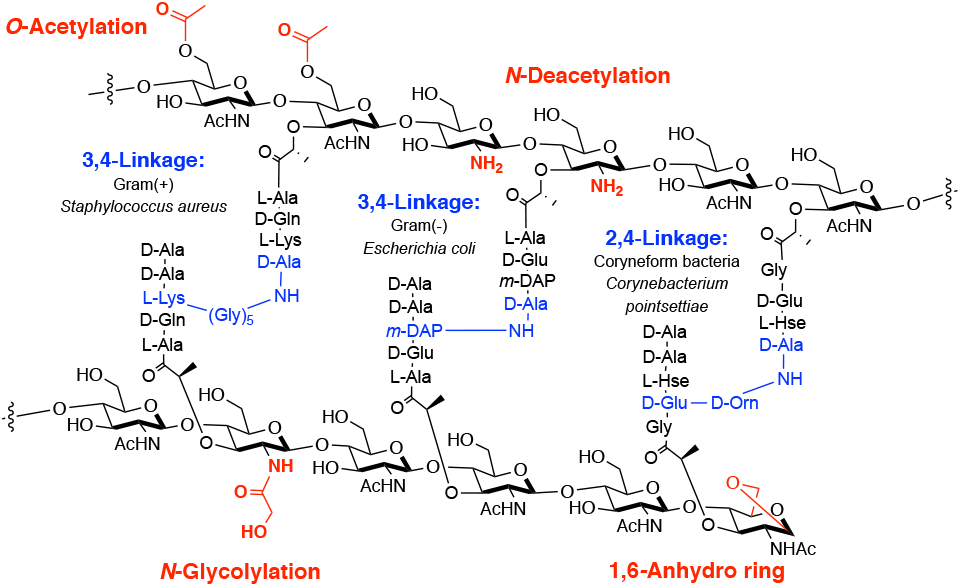
Peptidoglycan structure and modifications. Chemical functionality of both the peptide sidechain (blue) and carbohydrate backbone (red) in bacterial PG across species.

Variations in PG structure among bacterial species have led chemists to synthesize a variety of small molecule probes based on monomeric units of this structure. In particular, many of these building blocks are now commercially available. For example, *N-*acetyl muramyl dipeptide (MDP) is a representative small molecule PG mimic for “general” bacteria and has been shown to interact with NLRs, such as NOD2 and NLRP1, which are associated with a variety of diseases including Irritable Bowel Diseases (IBDs) and Vitiligo^40-42^ (Scheme 1). In addition, a “modified” bacterial PG fragment, *N*-glycolyl MDP (ng-MDP), derived from the hydroxylated product of *Mycobacterium*, has been heavily investigated due to *M. paratuberculosis’* direct link with CD^43^ (Scheme 1). Both “general” and “modified” PG fragments have been shown to stimulate a potent NOD2 dependent immune response^27^ and as such are the primary ligands of choice for immunologists due to the ligand synthetic simplicity and commercial availability (Scheme 1).

However, MDP and ng-MDP only represent a defined element of the PG fragment pool and fail to capture the major PG hydrolase degradation product, disaccharide muropeptides^44-45^. These disaccharide PG fragments are synthetically complex, limiting their accessibility and restricting their analysis in traditional immunostimulatory assays such as NFκB luciferase-reporter and ELISA screens^46-49^. Questions in the field surrounding the natural ligand of NLRs, like NOD2, as well as the biological influence of other *N*-acetyl muramic acid-containing PG fragments, still remain unanswered without an expanded cell wall library^50^. In this study, disaccharide PG-based fragments were identified from a gut microbe, *Lactobacillus acidophilus*^*51-52*^. Inspired by the generation of these PG products, a reliable and scalable synthetic route to several PG disaccharide fragments was implemented, leading to the first fully characterized *N*-acetylglucosamine *N-*acetylmuramic acid tripeptide (GMTP). These disaccharides were combined with a library of monosaccharide PG derivatives^53-54^ and screened for gene transcription activation, cytokine production and phosphorylation profiles using bone-marrow derived macrophages (BMDMs). Interestingly, GMTP is more potent activator for select gene transcriptional programs, many of which are related to IBD, when compared to their monosaccharide counterparts. Measuring cytokine production and downstream phosphorylation programs complement these genetic studies. Gene expression profiles comparing the disaccharide to the monosaccharide fragments, reveal a complex innate immune signaling pattern, validating a highly intricate molecular mechanism for sensing PG fragments.

## Results and Discussion

### Identification of Disaccharide PG Fragments from *L. acidophilus* Cultures

Previous exploration of Gram-negative and Gram-positive bacteria has led to the identification of several disaccharide PG fragments that activate an immune response^27, 49, 55-60^. In order to screen for biologically relevant fragments from Gram-positive bacteria residing in the gut microbiome, a lysozyme degradation assay of *L. acidophilus* was implemented (Scheme 2). *L. acidophilus* is a major commensal inhabitant of the human microbiota in intestinal, oral and vaginal tracts^61^. Since the 1970s, *L. acidophilus* has been commercially produced as a probiotic^52^ with reported therapeutic effects^62^. To screen this organism for PG small molecule production, *L. acidophilus* cultures were treated with lysozyme^63^, a muramidase found in high concentrations in both the human mouth and gut^64^. The lysed PG was then subjected to high resolution liquid chromatography mass spectrometry (HR-LCMS) analysis, and four disaccharide PG fragments were identified: *N*-acetylglucosamine *N-*acetylmuramic acid dipeptide (GMDP), *N*-acetylglucosamine *N-*acetylmuramic acid tripeptide (GMTP), *N*-acetylglucosamine *N-* acetylmuramic acid tetrapeptide (GMTTP), and *N*-acetylglucosamine *N-*acetylmuramic acid pentapeptide (GMPP) (Scheme 2). GMTP was observed in high abundance compared to other disaccharide fragments (Supplementary Information (SI) Table 2-Entry 2). This result is in agreement with previously published work in which GMDP, GMTP, GMTTP, and GMPP ^51, 57-59, 63, 65-69^ have been identified with variations at the third amino acid residue. However, lack of accessibility to these disaccharides, due to the complexity of chemical synthesis, has prevented a rigorous investigation of their immunological activity. This motivated the synthetic development of well-characterized disaccharides in larger and more accessible quantities.

**Scheme 2.**
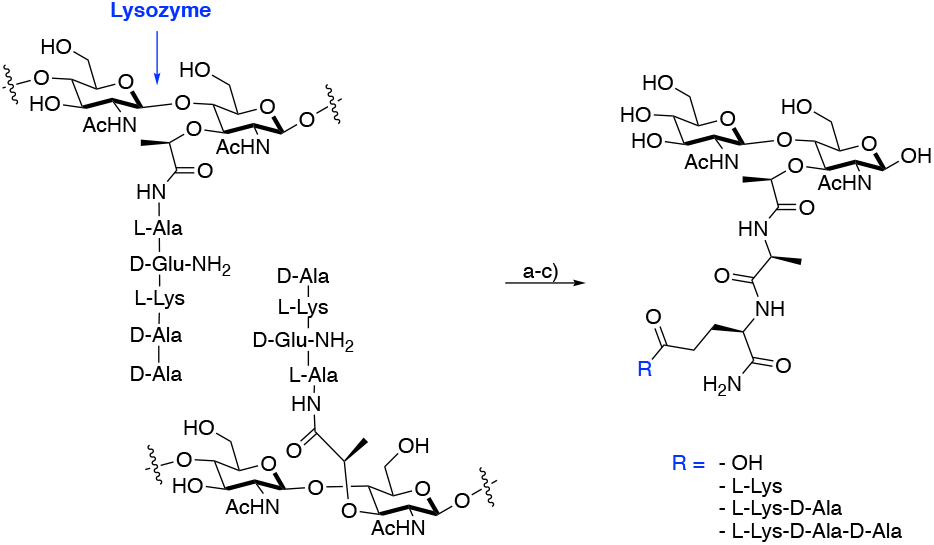
Lysozyme degradation assay for *L. acidophilus*. (a) Whole bacterial cells were treated with lysozyme, (b) lysed PG was subjected to a 3kDa spin filter, (c) PG fragmentation was analyzed by high resolution liquid chromatography mass spectrometry (HR-LCMS) and four fragments were identified (SI Table 2). Assay was replicated on three occasions. None of the identified masses were observed in the controls: media, wash buffer or enzymatic treatment buffer.

### Synthesis and Characterization of Disaccharide PG Fragments

A total synthesis using Schmidt glycosylation was implemented to obtain the protected β-1,4 linked intermediate **10** over 13 chemical transformations^57, 70-71^. From this intermediate, a modular strategy was utilized to access PG fragments **1-3** (Scheme 3). Compound **1** was first produced through the global deprotection of the acetyl protecting groups, as well as (Trimethylsilyl)ethyl Ester (TMSE), of compound **10**, followed by direct hydrogenation with 20% Pd(OH)_2_ to yield GMMP (**1**). Compound **2** was obtained through the deprotection of TMSE with 1N TBAF, followed by direct coupling of _D_-isolgn-OBn to yield intermediate **11**. Intermediate **11** was then de-acetylated using aqueous LiOH, followed by hydrogenation with 20% Pd(OH)_2_ to yield GMDP (**2**). Finally, the TMSE group was deprotected from **10** and directly coupled to the protected dipeptide, _D_-isolgn-_L_-Lys(Z)-OBzl, to yield **12**. Using a slightly modified synthesis, _D_-isolgn-_L_-Lys(Z)-OBzl was synthesized over two steps starting from commercially available H-Lys(Z)-OBzl and Boc-_D_-glutamic acid α-amide^53^. **12** was then deprotected in two subsequent steps to yield the final product GMTP (**3**). To ensure purity for biological testing, all final compounds were purified via reverse chromatography using a mass directed auto-purification system. From intermediate **10**, compounds **1-3** were obtained with an overall yield of 56% (2 steps), 9% (4 steps), and 21% (4 steps), respectively. Full NMR characterization and spectral data are presented for each synthetic compound, which has been previously unavailable to date, for synthetic fragment GMTP (**3**)^72^.

**Scheme 3:**
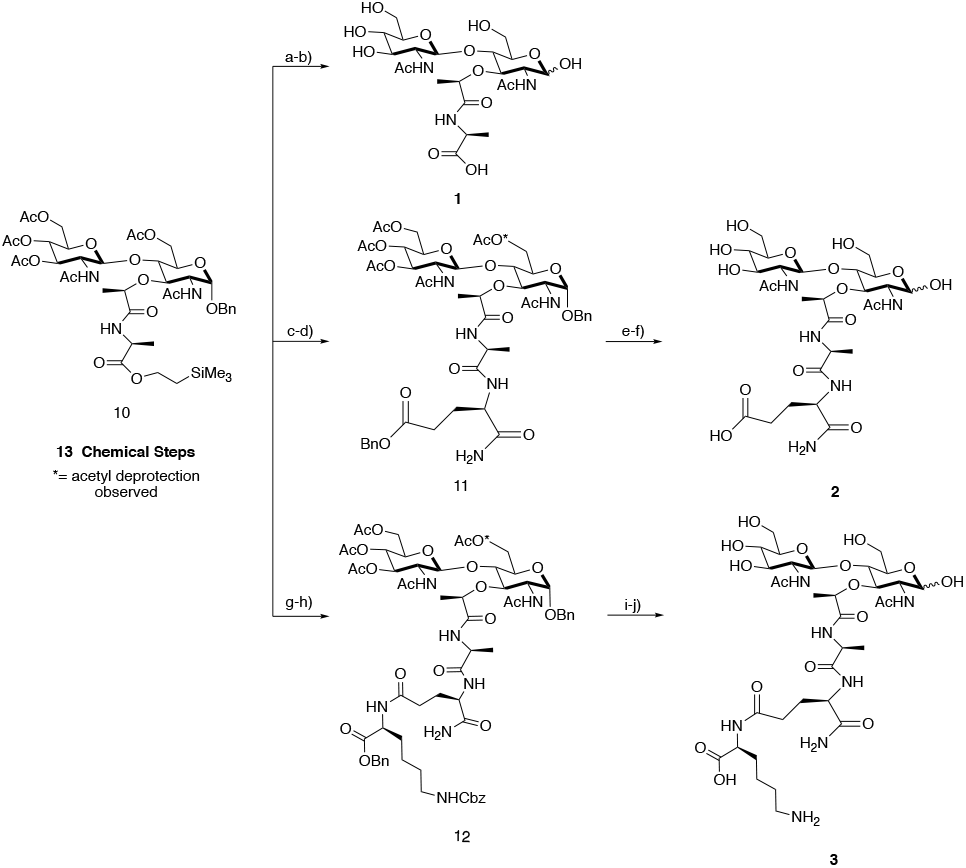
Synthesis of Disaccharides. Intermediate **10** was synthesized over 13 chemical steps. GMMP (**1**): (a) LiOH, ACN/H_2_O, (b) 20% Pd(OH)_2_, H_2_, THF/H_2_O yield 56%, over two steps. GMDP (**2**): (c) 1N TBAF in THF, (d) DIPEA, HBTU, HOBt, _D_-isogln-OBn, DMF yields 24%, over two steps. (e) LiOH, ACN/H_2_O, (f) 20% Pd(OH)_2_, H_2_, THF/H_2_O yield 37%, over two steps. GMTP (**3**): (g) 1N TBAF in THF, (h) DIPEA, HBTU, HOBt, _D_-isogln-_L_-Lys(Z)-OBzl, DMF yields 58%, over two steps. (i) LiOH, ACN/H_2_O, (j) 20% Pd(OH)_2_, H_2_, THF/H_2_O yield 37%, over two steps.

The assignment of GMTP (**3**) was confirmed utilizing a variety of 2D NMR experiments including ^1^H–^13^C HSQC-TOCSY, ^1^H–^1^H COSY, ^1^H–^13^C HSQC and ^1^H–^13^C HMBC (Figure 2, SI). Upon hydrogenation, the anomeric hydroxyl mutarotates into a mixture of two α/β isomeric species, with the α anomer being favored. Through detailed NMR experiments, the elucidation of the saccharide residues and peptide chains were obtained, confirming the structure and purity of the synthetic PG fragments.

**Figure 2:**
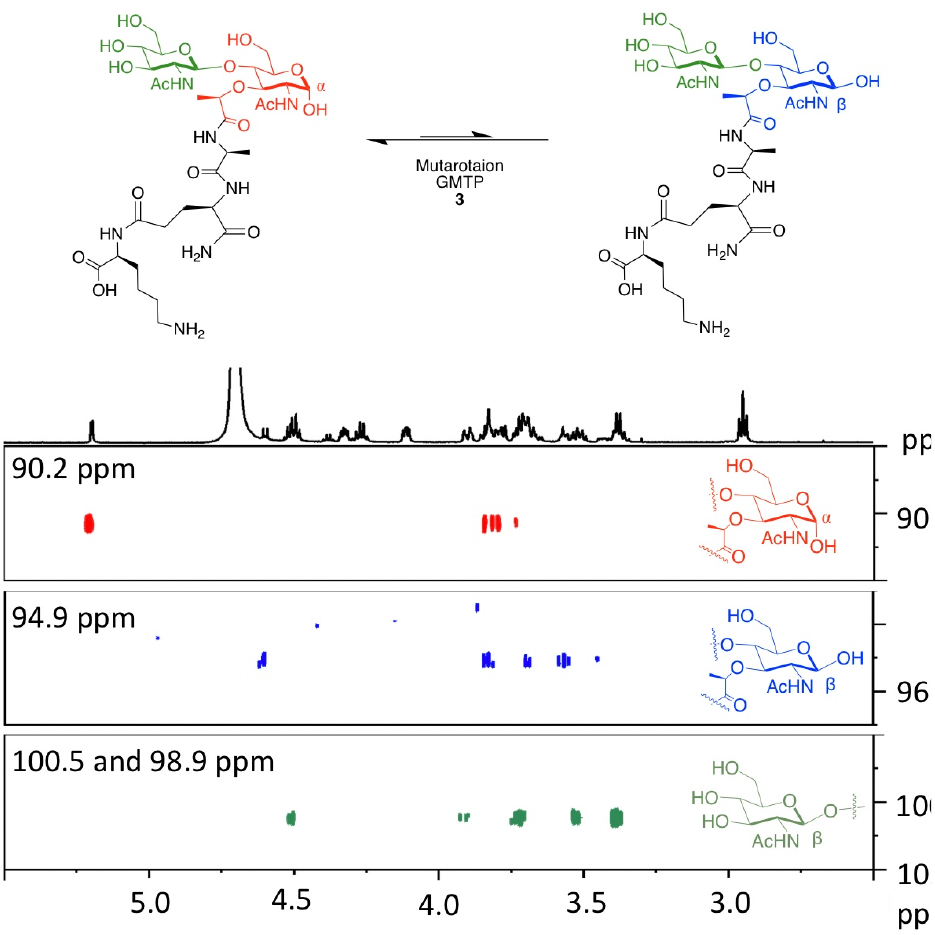
GMTP (3) ^1^H–^13^C HSQC-TOCSY. ^1^H–^13^C HSQC spectra of GMTP (**3**) with magnification of anomeric region for major and minor isomer carbohydrate ring spin systems. Spectra were recorded on a 600 mHz Bruker NMR at 298 K in D_2_O with settings: D9= 0.12, O1P= 3ppm, O2P= 60ppm, SW= 6ppm, 1SW= 100ppm, D1= 2sec, NS= 24.

### Genome-wide Transcriptional Analysis Reveals PG Disaccharide regulated Target Genes

With access to large quantities of compounds **1-3**, an investigation of the regulation programs in mouse bone-marrow derived macrophages (BMDMs) in response to both mono- and disaccharide PG fragments was implemented. To investigate gene regulation mediated by PG units, a qRT-PCR assay was first utilized to analyze the expression of immune response indicator genes. A subset of nine compounds containing both synthesized disaccharides **1-3** and six additional monosaccharides were used to stimulate macrophages (SI-Figure 1). In this initial investigation, target genes were selected based on previous RNA sequencing analysis with ng-MDP^73^. From this screen, the mRNA expression level of the tested genes increased significantly in BMDMs treated with GMTP compared to MDP and ng-MDP (Figure 3A and 3B, SI-Figure 2). Gene expression regulation was observed after 4 hours of stimulation with of 20 μM compound treatment (Figure 3A and 3B, SI-Figure 2). Importantly, this upregulation was not observed when GlcNAc was removed from the GMTP structure to yield the monosaccharide *N-*acetylmuramic acid tripeptide (MTP), demonstrating that the β-1,4 linked GlcNAc residue plays a critical role in the observed gene activation (SI-Figure 1 and 2). Additionally, the third amino acid, lysine, could play a critical role in generating a significant gene response as other disaccharide fragments such as GMMP (**1**) and GMDP (**2**) did not activate as robustly (Scheme 2, SI-Figure 2). Upon further investigation, GMTP was determined to activate *Tnf-*α and *Cxcl10* significantly more than MDP and ng-MDP (Figure 3B, SI Figure 3). *Il-1*β, and *Cox2* transcripts were also potently induced by GMTP, whereas MDP had no activation (Figure 3B). These results showcase for the first time a specific disaccharide PG unit capable of initiating gene expression programs differently than the previous monosaccharide PG standards, MDP and ng-MDP.

**Figure 3:**
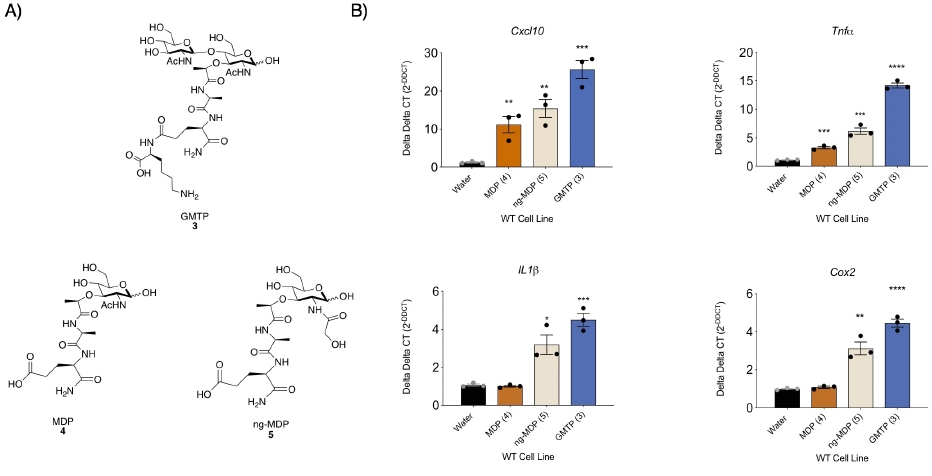
GMTP is a potent inducer of proinflammatory cytokines. A) Molecular structures of compounds GMTP (**3**), MDP (**4**) and ng-MDP (**5**). B) Gene expression statistical analysis of qRT-PCR data for *Tnf*α, *Il-1*β, *Cox2*, and *Cxcl10* in BMDMs treated with 20µM of (**3)**, (**4**), (**5**) or control (water) for 4 hours. Total RNA was harvested, and the expression levels of select genes were analyzed by qRT-PCR. Individual ΔΔCT values shown in SI-Table 1. Error bars indicate mean±SEM. Statistical significance was calculated using Student’s two-tailed t test. *P≤0.05, **P≤0.01, ***P≤0.001, and ****P≤0.0001 (n=3).

To more precisely understand the transcriptional programs upon PG fragment stimulation and measure the differences observed in the qRT-PCR analysis, whole genome RNA sequencing (RNAseq) study was performed on BMDMs treated for 18h with 20 µM GMTP, MDP, MTP, and ng-MDP (Figure 4A, SI-Figure 1). Gene regulation in unstimulated (water) BMDMs was used as a control. DESeq2 and Intensity Difference analysis^74-75^ revealed significantly regulated genes in each stimulation group which were combined for gene set enrichment and hierarchical clustering in Seqmonk (Supplementary Information 2 (SI2), Tables 1-8). Here we show GMTP has significantly enhanced immune stimulating capacity compared to MDP, MTP and ng-MDP (Figure 4A, SI2 Table 9). GMTP induced a unique gene expression signature distinct from MDP (Figure 4A, SI2 (Tables 1, 2, 5 and 6)). Hierarchical clustering (HC) analysis identified genes primarily induced by GMTP such as Acod1, Ass1, Il6, GM17300 and CX3CL (Figure 4A, Cluster 1). Transcriptional responses to GMTP and MDP similarly induced genes of Cluster 2 that included Slfn2, Arl5c, Oas2, Gm36161 and Tmem176a (Figure 4A). In contrast, expression of S100a4, Rpl39, Rpns27l, Crip1 and Wdr89 was specifically induced by MDP (and to a lesser extent ng-MDP) but not the other PG fragments (Figure 4A, Cluster 5, SI2 (Table 1-9)). MTP was less efficient in upregulating genes that were characteristically induced by GMTP in Cluster 1 consistent with the finding that the GlcNAc component of GMTP is critical for gene activation (SI2 (Tables 5 and 7)). ng-MDP, the “modified” PG fragment, induced a strong response of a cluster of genes LTF, LCN2, Ngp, Chil3, Mmp8, MMp9, CD177, S100a9, S100a8, Il2rb, Lck and Retnlg but not GMTP in Cluster 4 (SI2, (Tables 4 and 8)), indicating a monosaccharide specific expression pattern. Genes in Cluster 3 of the analysis where induced by GMTP, MDP and MTP (and not the modified PG, ng-MDP), representing a core, “general” PG response signature that includes IL1b, IFF7, Slfn5, Slfn8, Icam1, Ifit2, Ifit3 Oas3, Rsad2 Ifi209 and Tmem176b (Figure 4A). Overall, macrophages responded to different PGs with identifiable unique gene expression signatures.

**Figure 4.**
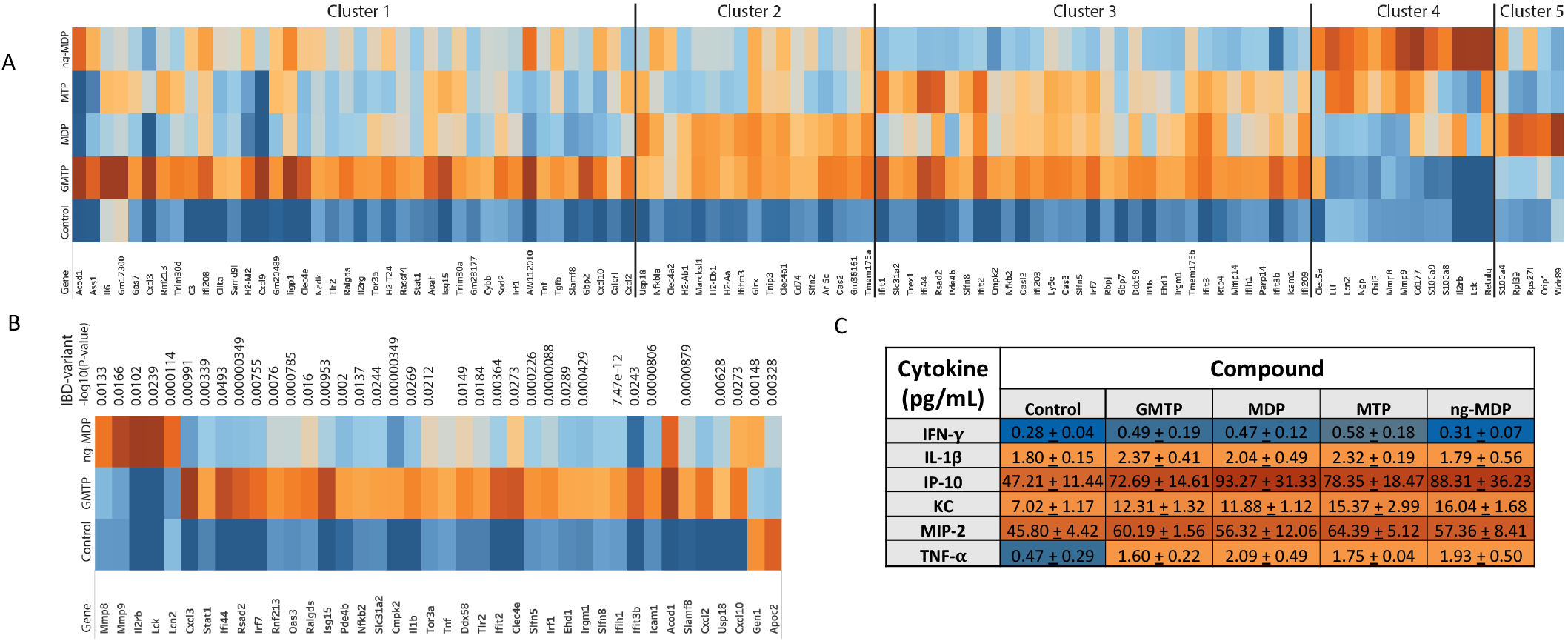
RNA sequencing analysis of the differential gene expression in BMDMs after treatment with PG fragments. BMDMs derived from WT mice were treated with PG fragments GMTP, MDP, MTP, or ng-MDP at 20 μM for 18h (n=3, biological replicates for each sample). Total RNA was subjected to RNA sequencing analysis (A & B). A) Heatmap of top genes differentially expressed (p < 0.01, FDR < 0.05, and logFC > 1) in BMDMs treated with water (control) or PG fragments GMTP, MDP, MTP, or ng-MDP. Hierarchical clustering separated genes into 5 clusters, dependent upon the PG activation pattern. Color scales represent upregulation (red) or downregulation (blue) of respective genes. (B) Heatmap showing all the top IBD gene sets that are upregulated (red) or downregulated (blue) differentially between GMTP and ng-MDP compared to the control (water). (C) Luminex analysis performed on supernatants derived from media culture from BMDM + 20 μM compound treatment for 18h. Data is represented as mean +/- SEM (n=3, biological replicates).

GMTP emerged from these experiments as a new and efficient activator of innate immune responses in macrophages. Remarkably, 31 genes that were identified by DEseq2 analysis^74-75^ as significantly induced by GMTP have genetic variants associated with either Crohn’s disease or ulcerative colitis (IBD exome browser, ibd.broadinstitute.org) (Figure 4B, SI2, Table 10). The relative expression of these IBD associated genes and the P values of the highest IBD associated variant are shown (Figure 4B, SI2 Table 10). While a few of these genes were also part of the ng-MDP induced pathways, we also identified IBD associated genes that characterized the response to ng-MDP including Mmp8, Mmp9, IL2rb, Lck and Lcn2 (Figure 4B, SI2 Table 10). These findings indicate that the pathway associated with the recognition of GMTP may play an import role in regulating mucosal immune responses.

An ELISA-Luminex based assay confirmed that IL-1β, CXCL10 (IP10), KC (IL8 homologue), MIP2, IFN-y and TNF-α protein expression was induced in the PG-fragment treated macrophages (Figure 4C). Together, these results confirmed that the synthesized peptidoglycan fragments were potent innate immune stimuli that were able to induce shared and fragment specific gene expression profiles that maybe able to elicit unique immune responses.

### Cellular Biochemical Characterization

To further extend this study from gene to protein level, we next analyzed the activation, i.e. phosphorylation, of common PG signaling pathways. Phosphorylation events are an essential component of the downstream production of cytokines and chemokines upon PG stimulation (Scheme 1)^73^. Through immunoblot analysis, Stat1, IRF5, cJun, and p65 NFκB phosphorylation was screened (Figure 5). Phosphorylation was observed for IRF5, cJun and P65-NFκB at 1h and 4h treatment with 20 uM MDP, ng*-*MDP and GMTP suggesting that all the three compounds were able to activate these signaling pathways. The stronger transcriptional activation of immunoregulatory genes observed for GMTP in combination with the RTPCR and RNAseq analysis (Figure 3B, 4A, and 4B), could be a result of combinatorial activation of these pathways, or involve yet uncharacterized transcriptional regulators. The stability or component recruitment of PRRs (Figure 1) into the subsequently forming signaling complexes may be distinct and responsible for the observed fragment specific transcriptional responses. Future studies probing the potential PRR(s) binding mechanism and complexes components are needed to further understand these PG-associated innate immune responses.

**Figure 5.**
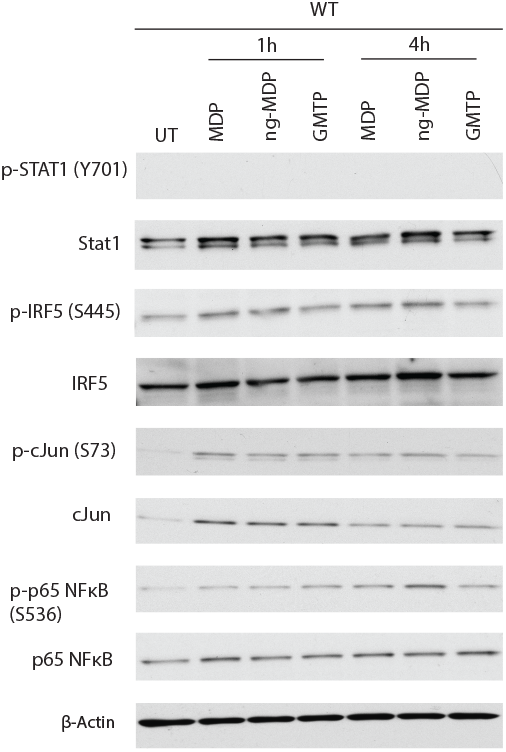
Induction of Stat1, IRF5, cJun, and p65 NFκB phosphorylation in BMDMs with 20 µM MDP, ng-MDP, GMTP and control (UT, water) isolated at 1 hour and 4 hours.

## Conclusion

Key lysozyme products of the Gram-positive commensal gut bacterium, *L. acidophilus*, were identified and confirmed. Corresponding disaccharide biologically relevant PG fragments were subsequently synthesized and fully characterized on scale. Whole genome RNA-sequencing reveal that GMTP was identified as a key disaccharide PG fragment that serves as significant activator of multiple gene transcriptional and cytokine programs (Figure 4A and 4B). From the biological assay results, it appears different muramyl peptides and glycans can tune the transcriptional activation of a variety of immunoregulatory genes through the combinatorial activation of distinct transcription factors such as NFκB, cJUN and IRF5^73^. Remarkably, GMTP treatment revealed a new gene cluster that is associated with IBD (Figure 4B). These results complement the sophisticated, yet limited reports in the field that use adjuvants other than MDP, such as GMTP-N-dipalmitoylpropylamide (DPG)^78^ and mifamurtide^79^, for potent stimulation of PRR receptors. The knowledge of these specific activation pathways will allow for more tunable adjuvants to be compiled.

This work shows that distinct PG fragments yield a variety of inflammatory gene patterns. Innate immune stimulation by bacterial peptidoglycan is much more complex than previously observed with the small synthetic PG mimics, muramyl dipeptide (MDP) and *N*-glycolyl MDP (ng-MDP) with supporting evidence emerging in the recent literature^69,73^. These distinct molecular signatures of pathogen and commensal bacteria could allow for PRRs to maintain homeostasis and regulate inflammation within the microbiome. A next obvious step forward from this work is to catalog all biologically relevant PG fragments. With these molecules readily available to the scientific community, harnessing the mechanisms, as well as structural details, responsible for this differential PG signaling paradigm will be essential in developing novel therapeutics to control inflammation.

## ASSOCIATED CONTENT

Supporting Information

Supplementary Information 1 (SI, pdf) includes supplementary figures and tables, biochemical methods, synthetic procedures and compound characterization. Supplementary Information 2 (SI2, .xls) includes RNAseq data tables 1-10.

## AUTHOR INFORMATION

Corresponding Authors

*

The authors declare no competing financial interest.

## AUTHOR CONTRIBUTION

The manuscript was written through contributions of all authors. All authors have given approval to the final version of the manuscript.

## ACKNOWLEDGMENTS

This work was supported by the Delaware COBRE program, with a grant from the National Institute of General Medical Sciences (NIGMS P20GM104316; C.L.G.) and the National Science Foundation (NSF 1554967). C.L.G. is a Pew Biomedical Scholar and Sloan Fellow and thanks the Pew and Sloan Foundations. HCR and CLG acknowledge GM138599 from the National Institutes of Health; HCR acknowledges AI11333, and DK068181 from the National Institutes of Health. K.M.L. and K.E.D. thank the University of Delaware for their support through the University Doctoral and Dissertation Fellowship Programs. K.E.D. and K.A.W. thank the NIH for support (5T32GM008550). Instrumentation support was provided by the Delaware COBRE and INBRE programs, supported by the National Institute of General Medical Sciences (P30 GM110758, P20 GM104316, and P20 GM103446). Thank you to Dr. PapaNii Asare-Okai for mass spectrometry and liquid chromatography support. A special thank you to Dr. Shi Bai for NMR support in obtaining the spectral data for 1D and 2D experiments.

## ABBREVIATIONS

(MDP): Muramyl dipeptide
(PAMPS): pathogen associated molecular patterns
(MAMPS): microbe associate molecular patterns
(PRRs): pattern recognition receptors
(TLRs): Toll-like receptors
(CLRs): C-type lectin receptors
(NLRs): NOD-like receptors,
(RLRs): RIG-I-like receptors
(ALRs): AIM2-like receptors
(PGBPs): Peptidoglycan binding proteins
(SLAMF): SLAM family
(OLRs): OAS-like receptors
(DC): dendritic cells
(PG): peptidoglycan
(GMDP): N-acetyl glucosamine muramyl dipeptide
(GMTP): N-acetyl glucosamine muramyl tripeptide
(GMTTP): N-acetyl glucosamine muramyl tetrapeptide
(GMPP): N-acetyl glucosamine muramyl pentapeptide
(NOD2): nucleotide-binding oligomerization domain-containing protein 2
(Mur-NAc): *N-*Acetyl Muramic Acid
(GlcNAc): *N-*Acetyl Glucosamine
(HSQC): Heteronuclear single quantum coherence spectroscopy
(HMBC): Heteronuclear Multiple Bond Correlation
(COSY): COrrelated SpectroscopY
(TOCSY): TOtal Correlated SpectroscopY

