## Supplemental Information 1 for "Bacterial Peptidoglycan Fragments Differentially Regulate Innate Immune Signaling"

### Table of Contents

I. **Materials and instrumentation**…………………………………………………………..3

II. **Synthesis of disaccharide fragments**…………………………………………………………4

III. **Bone marrow derived macrophages culture and treatment**……………………………...…9

IV. **Real-time quantitative-PCR**…………………………………………………………………...9

RNA isolation ……………………………………………………………………………..9
cDNA synthesis…………………………………………………………………………..…10
qRT-PCR and Primers …………………………………………………………………...…10
PG library (**SI Figure 1**) and results for qRT-PCR Initial Screen (**SI Figure 2**)……………..11-12
qRT-PCR GMTP vs. MDP 4 hour replicates (**SI Figure 3**)…………………………………...13
qRT-PCR Numerical **ΔΔCT** Data for Main Text Figure (**SI Table 1**)……………………...…14

V. **RNA Sequencing**……………………………………………………………………...……14

RNAseq protocols
Data Processing and Analysis

VI. **Biochemical Assays**………………………………………………………………………...15

Luminex protocol
Western blot analysis

VII. **Peptidoglycan Enzymatic Degradation and Mass Spectrometry Analysis**……………..……16

*Lactobacillus acidophilus* growth and enzymatic degradation
LC- Mass Spectrometry Method
Results and HRMS Spectra of Identified fragments (**SI Table 2**) ……………………………..…17

VIII. **NMR Spectra of Synthetic Compounds**……………………………………………………..20

IX. **HRMS Spectra of Synthetic Compounds**…………………………………………………...27

X. **References**…………………………………………………………………..…………29

#### I. Materials and Instrumentation

**Materials.**

All chemicals were purchased from Millipore-Sigma, ThermoFisher Scientific, ChemImpex and used without further purification unless otherwise noted. All solvents were reagent grade anhydrous purchased from Millipore-Sigma. Deuterated NMR solvents were purchased from Cambridge Isotope Laboratories. All compounds were purified on normal phase silica with flash chromatography unless otherwise noted. Total RNA was extracted using RNeasy Mini isolation kit (Qiagen). RNase-Free DNase Set (Qiagen) for DNase-digestion. QuantiTect Reverse Transcription Kit (Qiagen) was used for cDNA Synthesis. All primers were obtained from Eurofins Genomics LLC. Sybr SsoAdvanced Universal Green Supermix was obtained from Bio-Rad.

**Instrumentation.**

NMR spectra were recorded on either a Bruker AVIII 400 MHz or AV III 600 MHz spectrometers. High Resolution Mass Spectrometry (HRMS, ESI) data was obtained at the University of Delaware Mass Spectrometry Facility (Thermo Q-Exactive Orbitrap). Low-Resolution Mass Spectrometry (LRMS, ESI) were obtained using an ACQUITY UPLC H-Class/SQD2 or Shimadzu LCMS 2020 at the University of Delaware Mass Spectrometry Facility. Compounds purified on reverse phase were conducted on a Waters Autopurification System (2767 Sample Manager with HPLC and SQD2 MS with ESIA) with a SunFire Prep C18 OBD 5µM 18x100mm Column. PCR was performed on a 7300 Applied Biosystems RTPCR. RNA and DNA concentrations were obtained via nanodrop (Thermo Scientific). Whole genome RNA Sequencing data was generated on a Illumina NextSeq 550 system at the University of Delaware Sequencing Core. Luminex was performed at the University of Maryland Baltimore Cytokine Core using a Luminex MagPix reader.

#### II. Synthesis of disaccharide fragments

(A)

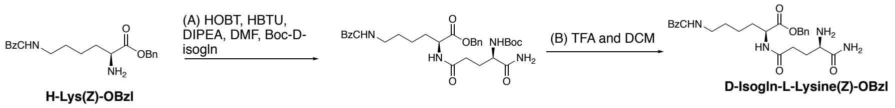


(B)

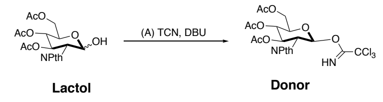


(C)

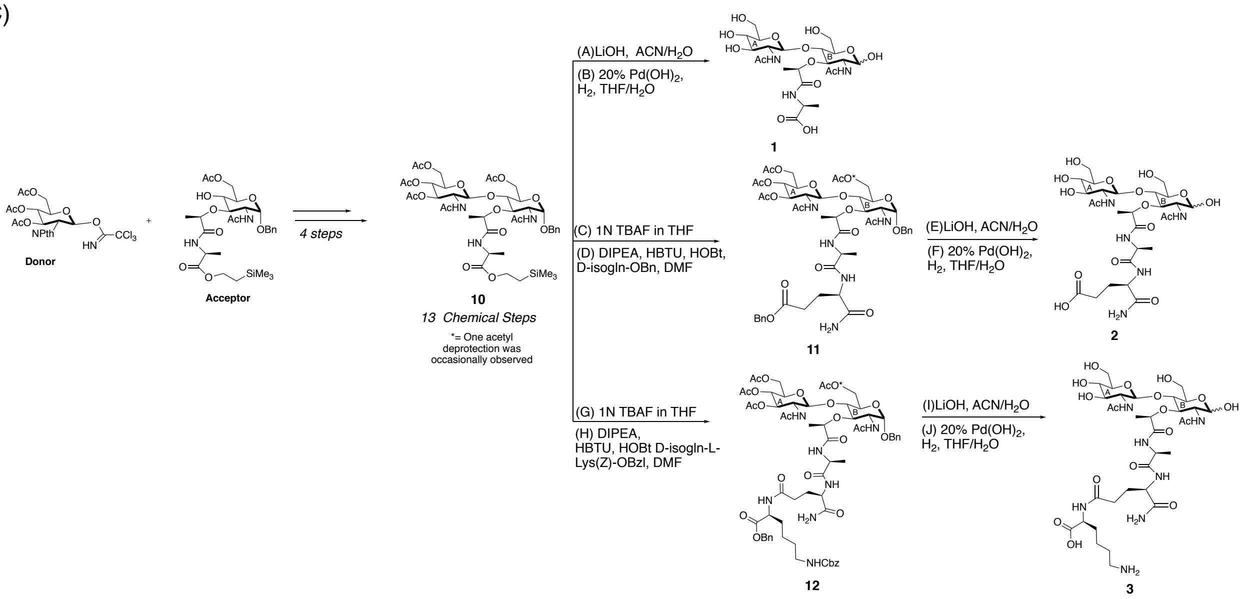


***D-isolgu- L-Lys(Z)-OBzl:*** Boc-D-Glu-$NH_2$ (0.040 g, 0.16 mmols, 1 eq), HBTU (0.073 g, 0.195 mmols, 1.2 eq), HOBt (0.026 g, 0.195 mmols, 1.2 eq) and DIPEA (0.11 mLs, 0.64 mmols, 4 eq) were combined in a reaction round bottom flask with 1.1 mLs of anhydrous DMF on ice for 10 min. L-Lys(Z)-OBzl (0.079 g, 0.16 mmols, 1.2 eq) was added. After 24 hours, the reaction was diluted with ethyl acetate. The organic layer was then washed with 1N HCl, saturated sodium bicarbonate and brine (the water layers were re-extracted if product was detected in any of the water layers). The organic layer was dried with sodium sulfate and condensed. The crude was purified via column chromatography with a slow gradient (0-5% MeOH in DCM) to yield a white solid (50mg, 50%). The protected product (0.07g, 0.118mmols, 1eq) was then dissolved in 0.4 mLs of anhydrous DCM and then 0.2 mLs of TFA was added. The reaction was left to stir for 1.5 hours. The product was then condensed resuspended in water and lyophilized to yield a white fluffy powder (71 mg, quantitative). NMR and mass spectrometry data matched previously reported data[1a].

***Acceptor*:** Synthesized according to precedence. NMR matched previously reported[2].

***Donor:*** Lactol (4.25 g, 9.77 mmols, 1 eq) was dissolved in 49 mLs of DCM then trichloroacetonitrile (19.5 mLs, 195 mmols, 20 eq) was added on ice followed by DBU (0.145 mLs, 0.977 mmols, 0.1 eq). The reaction stirred at 0˚C for 5 hours the reaction was then condensed and purified by column chromatography with 0-40% EtOAc/Hexane to yield a white solid (5.18 g, 92%). NMR matched previously reported[2d, 3].

***(S)-2-((R)-2-(((3R,4R,5S,6R)-3-acetamido-5-(((2S,3R,4R,5S,6R)-3-acetamido-4,5-dihydroxy-6-(hydroxymethyl)tetrahydro-2H-pyran-2-yl)oxy)-2-hydroxy-6-(hydroxymethyl)tetrahydro-2H-pyran-4-yl)oxy)propanamido)propanoic acid (compound 1):*** Compound **10**[2] (0.032 g, 0.0345 mmol, 1 eq) was dissolved in 1.7 mL acetonitrile. Then 1.7 mL of 0.1 M $LiOH_{(aq)}$ was add dropwise. The reaction stirred overnight and was partially complete. An additional 0.35 mL 0.1 M $LiOH_{(aq)}$ was added and after two hours another 0.35 mL of 0.1M LiOH (aq) was added and left to stir five and half hours. Once complete (M+1=658), the reaction was quenched with acetic acid (monitored with pH paper to ~7), filtered, condensed and placed on high vac overnight. The product was re-suspended in 0.79 mL of THF and then 2.45 mL of DI water was added along with 0.21 mL of acetic acid. 20% $Pd(OH)_2$ (12 mg, 0.0172 mmol, 0.5 eq) was added, and the flask was evacuated and filled with hydrogen three times. After stirring overnight, the reaction was confirmed as complete then filtered and condensed. The residue was purified via reverse phase SunFire Prep C18 column using an Autopure (0-20% acetonitrile in water with 0.1% Formic acid over 5 min) to yield a white powder (11 mg, 56%).α:β ratio 1:0.3 for carbohydrate B (anomeric center). α isomer is designated as major isomer, β is designated as minor isomer. NMR assignments based on 2D experiments (α major isomer is predominantly assigned). $^1H$ NMR (600 MHz, Deuterium Oxide) δ 5.16 (d, *J* = 3.4 Hz, 1H, $H_1B_{major}$), 4.59 (d, *J* = 7.4 Hz, 0.24H, $H_1B_{minor}$), 4.50 (d, *J* = 8.4 Hz, 1H, $H_1A_{major}$), 4.48 (d, *J* = 8.4 Hz, 0.40H, $H_1A_{minor}$), 4.44 (q, *J* = 6.6 Hz, 1H, C*H*-propionic$_{major}$), 4.38 (q, *J* = 6.5 Hz, 0.43H, C*H*-propionic$_{minor}$), 4.11 (qd, *J* = 7.3, 4.0 Hz, 1.59H, C*H*-alanine$_{major/minor}$), 3.92 – 3.88 (m, 1.49H, $H_6A'_{major}$, $H_6A'_{minor}$ ), 3.86 – 3.62 (m, 12.82H, $H_6B'_{minor}$, $H_2B_{major}$, $H_6B'_{major}$, $H_4B_{major}$, $H_4B_{minor}$, $H_6B''_{major}$, $H_6B_{minor}$, $H_6A''_{major}$, $H_6A''_{minor}$, $H_3B_{major}$, $H_2A_{major}$, $H_2A_{minor}$, $H_5B_{major}$, $H_2B_{minor}$), 3.52 (ddd, *J* = 10.5, 7.9, 6.2 Hz, 1.64H, $H_3A_{major}$, $H_3A_{minor}$), 3.45 (ddd, *J* = 9.8, 5.1, 2.1 Hz, 0.52H, $H_3B_{minor}$), 3.43 – 3.34 (m, 3.3H, $H_4A_{major}$, $H_5A_{major}$, $H_4A_{minor}$, $H_5A_{minor}$), 2.01 (s, 5H, N*H*Acetate$_2$A$_{major/minor}$), 1.94 (s, 3H, N*H*Acetate$_2$B$_{major}$), 1.92 (s, 1H, N*H*Acetate$_2$B$_{minor}$), 1.37 – 1.30 (m, 10H, C*H*$_3$-Alanine$_{major/minor}$, C*H$_3$*-Propionic$_{major/minor}$). $^{13}C$ NMR (151 MHz, $D_2O$) δ 174.70 (NH-Acetate$_2$A$_{major/minor}$), 174.64(NH-Acetate$_2$B$_{minor}$), 174.35(Carbonyl-Alanine$_{major/minor}$), 174.28 (NH-Acetate$_2$B$_{major}$), 173.60 (Propinic-Carbonyl$_{major/minor}$), 100.50 ($C_1A_{minor}$), 100.39 ($C_1A_{major}$), 95.27($C_1B_{minor}$), 90.44($C_1B_{major}$), 79.05 ($C_5B_{minor}$), 78.29(C-Propionic$_{minor}$), 78.03 (C-Propionic$_{major}$), 76.59($C_3B_{major}$), 76.19($C_5A_{minor}$), 76.16($C_5A_{major}$), 75.54($C_4B_{minor}$), 75.50($C_4B_{major}$), 75.17($C_3B_{minor}$), 73.62($C_3A_{major/minor}$), 70.99($C_5B_{major}$), 70.32($C_4A_{major}$), 70.30 ($C_4A_{minor}$), 61.12($C_6A_{major}$), 61.11($C_6A_{minor}$), 59.93($C_6B_{minor}$), 59.83($C_6B_{major}$), 56.09($C_2A_{major}$), 56.02 ($C_2A_{minor}$), 55.41 ($C_2B_{minor}$), 53.41 ($C_2B_{major}$), 51.20 (C-Alanine$_{major/minor[based\ on\ HSQC]}$) 22.45($C_2$B-Acetate$_{minor}$), 22.16 ($C_2$B-Acetate$_{major}$), 22.13 ($C_2$A-Acetate$_{major}$), 17.78($CH_3$-Propionic$_{major}$), 17.27 ($CH_3$-Propionic$_{minor}$), 17.03 ($CH_3$-Alanine$_{major}$), 16.95($CH_3$-Alanine$_{minor}$). HRMS m/z: $[M+H]^+$ calcd $C_{22}H_{38}N_3O_{14}^+$ 568.23483; Found 568.23615.

***(2R,3S,4R,5R,6S)-5-acetamido-6-(((2R,3S,4R,5R,6S)-5-acetamido-4-(((R)-1-(((S)-1-(((R)-1-amino-5-(benzyloxy)-1,5-dioxopentan-2-yl)amino)-1-oxopropan-2-yl)amino)-1-oxopropan-2-yl)oxy)-6-(benzyloxy)-2-(hydroxymethyl)tetrahydro-2H-pyran-3-yl)oxy)-2-(acetoxymethyl)tetrahydro-2H-pyran-3,4-diyl diacetate (compound 11):*** Compound **10**[2] (0.180 g, 0.19 mmol, 1 eq) was dissolved in 4 mL of anhydrous THF. Then 1M TBAF in THF (0.77 mL, 0.77 mmol, 4 eq) was added. The reaction was partially complete in 3 hours, so an additional 0.3 mL (0.3 mmol, 2 eq) of 1M TBAF in THF was added. Reaction was done in an hour and condensed. The residue was dissolved in EtOAc and washed 2 x 1N HCl (water layer was checked for any residual product and re-extracted). The organic layer was dried, condensed and placed under high vacuum for one hour. A short column (2-7% EtOH in DCM) was run to remove residual TBAF and the product was used in the subsequent reaction (53%= -acetate product, 8%= product identified via mass spec).The major product (0.08 g, 0.101 mmol, 1 eq) and *D*-isoglutamine-benzylester HCl (0.037 g, 0.137 mmol, 1.2 eq) was then dissolved in 0.685 mL of DMF on ice. Then HOBt (0.0186 g, 0.137 mmol, 1.2 eq) and HBTU (0.052 g, 0.137 mmol, 1.2 eq) were added and stirred on ice for 3 minutes. DIPEA (0.079 mL, 0.46 mmol, 4 eq) was added dropwise on ice. The reaction then was warmed to room temperature and stirred till complete (1.5 hours). The reaction was condensed and diluted with EtOAc. The organic layer was washed 1N HCl, saturated bicarbonate, and brine. The organic layer was dried, condensed and purified via column chromatography (0-10% MeOH in DCM) to yield a white solid (product (-acetate)= 46 mg, 24% over two steps). Note: During reactions one acetate was deprotected and reported as the final product. $^{1}$H NMR (400 MHz, DMSO-$d_6$) δ 8.47 (d, $J$ = 6.1 Hz, 1H, N$H_B$Acetate), 8.28 (d, $J$ = 8.2 Hz, 1H, N*H*-D-isoglutamine), 8.09 (dd, $J$ = 19.8, 8.0 Hz, 2H, N$H_A$Acetate), 7.41 – 7.20 (m, 10H, $C_1$B-OBn, D-isoglutamine-OBn), 5.25 (t, $J$ = 9.9 Hz, 1H, $H_3$A), 5.08 (s, 2H, D-isoglutamine-C$H_2$Bn), 4.96 – 4.85 (m, 2H, $H_1$B, $H_4$A), 4.82 – 4.68 (m, 2H, $H_1$A, $H_6$B-OH), 4.62 (d, $J$ = 12.8 Hz, 1H, C$H_2$OBn$_B$'), 4.53 – 4.35 (m, 3H, C$H_2$OBn$_B$'', C*H*-propionic, C*H*-Alanine), 4.27 – 4.13 (m, 2H, C*H*-D-isoglutmaine, $H_6$A'), 3.94 (d, $J$ = 11.9 Hz, 1H, $H_6$A"), 3.84 – 3.51 (m, 6H, $H_2$A, $H_5$A, $H_2$B, $H_4$B, $H_6$B'"), 3.50 – 3.43 (m, 2H, $H_5$B, $H_3$B), 2.36 (t, $J$ = 8.3 Hz, 2H, C$H_2$-D-isoglutamine), 2.04 (C$H_2$-D-isoglutamine') 1.96 (d, $J$ = 4.4 Hz, 6H, AcetateH$_{4,6}$A), 1.92 (s, 3H, AcetateH$_3$A), 1.80 (s, 4H, N*H*AcetateA, C$H_2$-D-isoglutamine'"), 1.75 (s, 3H, N*H*AcetateB), 1.23 (dd, $J$ = 26.0, 6.8 Hz, 6H, C$H_3$-Alanine, C$H_3$-Propionic). $^{13}$C NMR (101 MHz, DMSO) δ 174.16 (propionic-Carbonyl), 173.37 (D-isolglutamine-carbonyl), 172.59 (Alanine-carbonyl), 172.51 (D-isolglutamine-carbonyl), 170.61 ($C_6$A-Carbonyl), 170.05 (NHAcetate-Caronyl), 170.03 (NHAcetate-Carbonyl), 170.02 ($C_3$A-Carbonyl), 169.79 ($C_4$A-Carbonyl) 138.09 (Aromatic), 136.60 (Aromatice), 128.90 (Aromatic), 128.68 (Aromatic), 128.47 (Aromatic), 128.38 (Aromatic), 127.92 (Aromatic), 127.83 (Aromatic), 99.68 ($C_1$A), 95.95 ($C_1$B), 76.41 (CH-Propionic), 76.23 ($C_4$B), 75.72 ($C_3$B), 72.63 ($C_3$A), 72.13 ($C_5$B), 70.65 ($C_5$A), 68.86 ($C_4$A), 68.64 ($C_1$B-*C*H$_2$OBn), 65.96 (D-isoglutamine-*C*H$_2$OBn), 62.16 ($C_6$A), 59.46 ($C_6$B), 54.62 ($C_2$A), 54.16 ($C_2$B), 51.96 (*C*H-D-isoglutamine), 48.56 (*C*H-Alanine), 30.44 (*C*H$_2$D-isolglutamine), 27.49 (*C*H$_2$D-isolglutamine), 23.11 (Acetate-*C*H$_3$), 23.05 (Acetate-*C*H$_3$), 20.89 (Acetate-*C*H$_3$), 20.85 (Acetate-*C*H$_3$), 20.78 (Acetate-*C*H$_3$), 19.32 (*C*H$_3$-Alanine), 19.21 (*C*H$_3$-Propionic). Note: 1,4-Beta-linkage was confirmed via HMBC, $C_1$A coupled to $H_4$B and $C_4$A coupled to $H_1$B. Major product (-OAc). HRMS m/z: [M+H-OAc]$^+$ calcd $C_{47}H_{64}N_5O_{19}^+$ 1002.41900; Found 1002.42053.

***(R)-4-((S)-2-((R)-2-(((3R,4R,5S,6R)-3-acetamido-5-(((2S,3R,4R,5S,6R)-3-acetamido-4,5-dihydroxy-6-(hydroxymethyl)tetrahydro-2H-pyran-2-yl)oxy)-2-hydroxy-6-(hydroxymethyl)tetrahydro-2H-pyran-4-yl)oxy)propanamido)propanamido)-5-amino-5-oxopentanoic acid (compound 2):*** Mono-deacetylated compound **11** (0.05 g, 0.050 mmol, 1 eq) was dissolved in 2.5 mL of acetonitrile, and 2.5 mL of 0.1M LiOH$_{(aq)}$ was added slowly. The reaction was stirred for 2 hours and was quenched with acetic acid (monitored with pH paper to ~7), filtered and condensed (Mass spec, M+23= 808, de-acetylated and de-benzyl ester intermediate) and subsequently used in the next step. The residue was then suspended in 1.2 mL THF then 3.75 mL DI water and 0.3 mL of acetic acid was added. 20% Pd(OH)$_2$ (17.5 mg, 0.025 mmol, 0.5 eq) was added and the flask was evacuated and filled with hydrogen three times. After

4 hours, another portion of 20% $Pd(OH)_2$ (17.5 mg, 0.025 mmol, 0.5 eq) was added, and the flask was evacuated and filled with hydrogen three times. The reaction stirred for an additional 24 hours under hydrogen until all starting material consumed as observed by LC-MS. The reaction was filtered and evaporated under reduced pressure. The residue was purified via reverse phase SunFire Prep C18 column using an Autopure (0-30% Acetonitrile in water over 5 min with 0.1% formic acid at 20mL/min) to yield a white powder (12.7 mg, 37%). α:β ratio 1:0.2 for Carbohydrate B anomeric center. α isomer is designated as major isomer, β is designated as minor isomer. NMR assignments based on 2D experiments (α major isomer is predominantly assigned). 1H NMR (600 MHz, Deuterium Oxide) δ 5.20 (d, J = 3.6 Hz, 1H, $H_1B_{major}$), 4.62 (d, J = 8.5 Hz, 0.19H, $H_1B_{minor}$), 4.58 – 4.48 (m, 2.12H, $H_1A_{major/minor}$, C*H*-propionic$_{major}$), 4.42 (q, J = 6.6 Hz,0.14H, , C*H*-propionic$_{minor}$), 4.29 (td, J = 21.8, 19.2, 11.8 Hz, 3.10H, C*H*-alanine$_{major/minor}$, C*H*-D-isoglu$_{major/minor}$), 3.92 (d, J = 12.3 Hz, 1.8H, $H_6A'_{major/minor}$), 3.85 (q, J = 7.9 Hz, 3H, $H_3B_{major}$, $H_4B_{minor}$), 3.82 – 3.78 (m, 2H, $H_2B_{major}$, $H_6B'_{major/minor}$), 3.76 – 3.65 (m, 8H, $H_2A_{major/minor}$, $H_4B_{major}$, $H_6B'_{major/minor}$, $H_6A'_{major/minor}$, $H_2B_{minor}$), 3.55 (td, J = 18.8, 17.8, 8.8 Hz, 2H, $H_3A_{major/minor}$, $H_3B_{minor}$), 3.43 – 3.37 (m, 4H, $H_4A_{major/minor}$, $H_5A_{major/minor}$, $H_5B_{major/minor}$), 2.33 – 2.20 (m, 4H, C*H*$_2$-D-isoglutamine), 2.1 (m, 1.3H, C*H*$_2$'-D-isoglutamine) 2.03 (s, 4H, NH-Acetate$_{major/minor}$), 1.99 – 1.88 (m, 7H, NH-Acetate$_{major/minor}$, C*H*$_2$''-D-isoglutamine), 1.42 (dd, J = 7.3, 2.0 Hz, 5H, $CH_3$-Alanine$_{major/minor}$), 1.37 (dt, J = 10.3, 6.0 Hz, 5H, $CH_3$-Propionic$_{major/minor}$). 13C NMR (151 MHz, D2O) δ 178.46 (carbonyl), 175.62 (carbonyl), 174.75 (carbonyl), 174.61 (carbonyl), 174.22 (carbonyl), 100.50 ($C_1A_{major/minor}$), 95.19 ($C_1B_{minor}$), 90.38($C_1B_{major}$), 78.0 (CH-Propionic$_{major/minor}$), 76.56 ($C_4B_{major}$), 76.18 ($C_5A_{major/minor}$), 75.56($C_5B_{minor}$), 75.32($C_4B_{minor}$), 73.69($C_3A_{major/minor}$), 71.21 ($C_3B_{major}$), 70.43 ($C_4A_{major/minor}$), 61.26 ($C_6A_{major/minor}$), 61.23 ($C_3B_{minor}$?), 60.11 ($C_6B_{minor}$), 59.91 ($C_6B_{major}$), 56.20 ($C_2A_{major}$), 56.13 ($C_2A_{minor}$), 55.80 ($C_2B_{minor}$), 53.73 ($C_2B_{major}$), 50.06 (CH-Ala$_{major/minor}$), 49.84 (CH-D-isolgutamine$_{major/minor}$), 31.74 ($CH_2$-D-isoglutamine$_{major/minor}$), 28.06 ($CH_2$-D-isoglutamine$_{major/minor}$), 22.4 (NH-Acetates-$CH_{3major/minor}$), 18.30 ($CH_3$-propionic$_{major/minor}$), 17.07 ($CH_3$-alanine$_{major/minor}$). HRMS m/z: $[M+H]^+$ calcd $C_{27}H_{46}N_5O_{16}^+$ 696.29341; Found 696.29087.

***(2R,3S,4R,5R,6S)-5-acetamido-6-(((2R,3S,4R,5R,6S)-5-acetamido-6-(benzyloxy)-4-(((9S,14R,17S,20R)-9-((benzyloxy)carbonyl)-14-carbamoyl-17-methyl-3,11,16,19-tetraoxo-1-phenyl-2-oxa-4,10,15,18-tetraazahenicosan-20-yl)oxy)-2-(hydroxymethyl)tetrahydro-2H-pyran-3-yl)oxy)-2-(acetoxymethyl)tetrahydro-2H-pyran-3,4-diyl diacetate (Compound 12):*** Compound **10**[2] (0.086 g, 0.093 mmol, 1eq) was dissolved in 2.16 mL of anhydrous THF. Then 0.4 mL of 1M TBAF in THF was added. After 3 hours, the reaction was stopped and condensed. The residue was then dissolved in 10% MeOH/EtOAc the organic layer was then washed 2x 1N HCl (water layer was checked for any residual product), dried and condensed and directly used in next reaction. The dry product was dissolved in 1.0 mL of DMF. D-isoglu-L-Lys(Z)-OBzl (0.072 g, 0.11 mmol, 1.2 eq), HOBt (0.015 g, 0.11 mmol, 1.2 eq), and HBTU (0.042 g, 0.11 mmol, 1.2 eq) was added at 0˚C. After 15min, DIPEA (0.064 mL, 0.371 mmol, 4.0 eq) was added then the reaction was brought to room temperature until complete. After 4 hours, the reaction was stopped and condensed. The residue was then dissolved in 10%MeOH/EtOAc and washed with 1N HCl, saturated bicarbonate and brine (water layer was checked for product). The organic layer was dried and condensed. The crude was then purified via column chromatography 0-7% MeOH in DCM (major product (-acetate)= 52 mg, 44%; minor product (no-deacetylation)= 17 mg, 14%. Total yield= 58% over two steps). Note: during reactions one acetate was deprotected and assigned as the major product. Major product NMR assignment (we note slight impurity of dipeptide): 1H NMR (600 MHz, DMSO-d6) δ 8.44 (d, J = 6.1 Hz, 1H, NH), 8.38 (d, J = 7.2 Hz, 1H, NH), 8.24 (t, J = 8.0 Hz, 3H, NH), 8.08 (d, J = 8.8 Hz, 1H, NH), 8.02 (d, J = 7.1 Hz, 1H, NH), 7.70 (s, 1H, NH), 7.66 (s, 1H, NH), 7.40 – 7.23 (m, 35H, Benzyl Protons), 7.21 (q, J = 5.7 Hz, 3H), 7.02 (d, J = 8.4 Hz, 1H, NH), 5.27 (dd, J = 12.0, 7.9 Hz, 1H, $H_3$A), 5.12 – 5.06 (m, 2H, Benzylester $CH_2$ ), 5.00 (s, 2H, CBz $CH_2$), 4.92 (d, 1H, $H_1$B), 4.89 (t, 1H, $H_4$A), 4.79 (d, J = 8.2 Hz, 1H, $H_1$A) , 4.67 (q, J = 7.9, 6.9 Hz, 1H, C6-OH), 4.61 (t, J = 11.2 Hz, 1H, $CH_2$OBn'), 4.50 – 4.36 (m, 3H, $CH_2$OBn'', CH-ala, CH-prop), 4.28 – 4.16 (m, 6H, $H_6$A, CH-lys), 4.01 – 3.92 (m, 2H, $H_6$A'', CH-D-isogln), 3.75 (t, J = 9.5 Hz, 2H, $H_4$B, $H_5$A), 3.68 (q, J = 9.0 Hz, 2H, $H_2$A, $H_6$B'), 3.63 – 3.51 (m, 2H, $H_2$B, $H_6$B''), 3.48 – 3.37 (m, 2H, $H_3$B, $H_5$B), 3.00 – 2.92 (m, 13H, ε-$CH_2$-Lys), 2.31 (t, J = 7.6 Hz, 2H, $CH_2$-D-isogln), 2.19 – 2.13 (m, 3H, $CH_2$-D-isogln'), 1.95 (d, J = 5.1 Hz, 8H, Acetate-$CH_3$x2, $CH_2$-D-isolgn''), 1.91 (s, 4H, Acetate-$CH_3$), 1.85 – 1.77 (m, 4H, Acetate-$CH_3$), 1.75 (d, J = 3.3 Hz, 3H, Acetate-$CH_3$), 1.69 (tt, J = 13.4, 7.2 Hz, 1H, β-$CH_2$-Lys'), 1.60 (dt, J = 14.2, 7.6 Hz, 2H, β-$CH_2$-Lys''), 1.37 (qd, J = 11.7, 9.8, 5.9 Hz, 6H, δ-$CH_2$-Lys), 1.27 (d, J = 6.7 Hz, 6H, $CH_3$-propionic, γ-$CH_2$-Lys), 1.22 (d, J = 6.9 Hz, 4H, $CH_3$-Ala). 13C NMR (151 MHz, DMSO) δ 174.16 (carbonyl), 173.54 (carbonyl), 172.66 (carbonyl), 172.58 (carbonyl), 172.50 (carbonyl), 172.36(carbonyl), 172.34 (carbonyl), 170.55 (carbonyl), 170.00 (carbonyl), 169.98 (carbonyl), 169.93 (carbonyl), 169.77 (carbonyl), 161.80 (carbonyl), 156.56(carbonyl), 156.54 (carbonyl), 138.13 (aromatic), 137.74 (aromatic), 137.72 (aromatic), 136.48 (aromatic), 136.43 (aromatic), 128.90 (aromatic), 128.88 (aromatic), 128.81 (aromatic), 128.80 (aromatic), 128.72 (aromatic), 128.68 (aromatic), 128.52 (aromatic), 128.46 (aromatic), 128.23 (aromatic), 128.20 (aromatic), 128.17 (aromatic), 127.93 (aromatic), 127.88 (aromatic), 99.70 ($C_1$A), 95.98 ($C_1$B), 76.39 (CH-Propionic), 76.29 ($C_4$B), 75.76 ($C_3$B), 72.67 ($C_3$A), 72.19 ($C_5$B), 70.72 ($C_5$A), 69.01 ($C_4$A), 68.71 ($CH_2$OBn), 66.32 (Benzylester-$CH_2$), 65.59 (CBz-$CH_2$), 62.22 ($C_6$A), 59.56 ($C_6$B), 57.41 (CH-D-isolgn), 54.73 ($C_2$A), 54.24 ($C_2$B), 52.6 (CH-Lys), 48.51 (CH-Ala), 31.95 ($CH_2$-D-isolgn''), 31.16 ($CH_2$-D-isolgn'), 30.92 ($CH_2$-β-Lys), 30.89 ($CH_2$-δ-Lys), 28.37 (residual dipeptide amine), 28.04 (residual dipeptide amine), 23.14(Acetate-$CH_3$, $CH_2$-γ-Lys), 23.07(Acetate-$CH_3$), 20.89(Acetate-$CH_3$), 20.84(Acetate-$CH_3$), 20.78 (Acetate-$CH_3$), 19.43 ($CH_3$-Prop), 19.28 ($CH_3$-Ala). Major product (-OAc). HRMS m/z: $[M+H-OAc]^+$ calcd $C_{61}H_{82}O_{22}N_7^+$ 1264.55074; Found 1264.55087.

***(S)-2-((R)-4-((S)-2-((R)-2-(((3R,4R,5S,6R)-3-acetamido-5-(((2S,3R,4R,5S,6R)-3-acetamido-4,5-dihydroxy-6-(hydroxymethyl)tetrahydro-2H-pyran-2-yl)oxy)-2-hydroxy-6-(hydroxymethyl)tetrahydro-2H-pyran-4-yl)oxy)propanamido)propanamido)-5-amino-5-oxopentanamido)-6-aminohexanoic acid (Compound 3)***: Mono-deacetylated compound **12** (0.017 g, 0.013 mmol, 1 eq) was dissolved in 0.65 mL of acetonitrile and 0.65 mL of 0.1M LiOH (aq). The reaction was complete in 2 hours and AcOH was added to stop the reaction (monitored with pH paper to ~7), filtered and condensed (Mass spec, M+23= 1070, de-acetylated and de-benzyl ester intermediate). The dry product was dissolved in 0.3 mL of THF, 0.1 mL of acetic acid and 1 mL of water. 20% $Pd(OH)_2$ (5 mg, 0.0065 mmol, 0.5 eq) was added and the flask was evacuated three times and filled with hydrogen. After 20 hours, reaction was gaged as incomplete and another portion of 20% $Pd(OH)_2$ (5 mg, 0.0065 mmol, 0.5 eq) was added. After 8 hours, reaction was still not done and an additional 20% $Pd(OH)_2$ (5 mg, 0.0065 mmol, 0.5 eq) was added. The reaction was left overnight. The reaction was then filtered and condensed. The crude product was purified by reverse-phase chromatography using an SunFire Prep C18 (0-7% acetonitrile in water with 0.1% formic acid over 4 minutes at 20mL/min) to yield as white solid

(4.1 mg, 37%). α:β ratio ratio 1:0.5 for Carbohydrate B anomeric center. α isomer is designated as major isomer, β is designated as minor isomer. NMR assignments based on 2D experiments (α major isomer was predominately assigned). $^{1}$H NMR (600 MHz, Deuterium Oxide) δ 5.20 (d, $J$ = 3.4 Hz, 1H, $H_1B_{major}$), 4.60 (d, $J$ = 8.4 Hz, 0.5H, $H_1B_{minor}$), 4.50 (dt, $J$ = 9.5, 7.3 Hz, 2H, $H_1A$, CH-$Prop_{major}$), 4.42 – 4.32 (m, 2H, CH-D-iso, CH-$Prop_{minor}$), 4.27 (q, $J$ = 7.5 Hz, 2H, CH-$Ala_{major/minor}$), 4.12 (dd, $J$ = 8.3, 5.2 Hz, 1H, CH-$Lys_{major/minor}$), 3.93 – 3.87 (m, 1H, $C_6A'_{major}$), 3.87 – 3.76 (m, 4H, $H_2B_{major}$, $H_4B_{major}$, $H_5B_{major}$, $H_6B'_{major}$), 3.76 – 3.64 (m, 4H, $C_6A''_{major}$, $C_2A_{major}$, $C_3B_{major}$, $C_6B_{major}$), 3.60 – 3.48 (m, 1H, $H_5A_{major}$), 3.38 (q, $J$ = 8.7, 7.3 Hz, 3H, $H_4A_{major}$, $H_5A_{major}$), 2.95 (t, $J$ = 7.5 Hz, 2H, Lys-$CH_2\varepsilon_{major/minor}$), 2.36 (td, $J$ = 7.8, 2.7 Hz, 2H, D-isogln-$CH_2\gamma_{major/minor}$), 2.17 – 2.08 (m, 1H, D-isogln-$CH_2\beta_{major/minor}$), 2.01 (s, 3, Acetate-$CH_3$-$A_{major/minor}$), 2.00 – 1.95 (m, 1H D-isogln-$CH_2\beta_{major/minor}$), 1.92 (d, $J$ = 10.6 Hz, 3H, Aceate-$CH_3$-$B_{major/minor}$), 1.77 (tt, $J$ = 13.5, 7.1 Hz, 1H, Lys-$CH_2\beta_{major/minor}$), 1.64 (qd, $J$ = 12.3, 9.9, 6.8 Hz, 3H, , Lys-$CH_2\gamma_{major/minor}$, Lys-$CH_2\beta_{major/minor}$), 1.42 – 1.31 (m, 8H, $CH_3$-$Ala_{major/minor}$, $CH_3$-$Prop_{major/minor}$, Lys-$CH_2\delta_{major/minor}$). $^{13}$C NMR (101 MHz, $D_2O$) δ 178.92 (Carbonyl), 175.93 (Carbonyl), 175.53 (Carbonyl), 175.10 (Carbonyl), 175.00 (Carbonyl), 174.91 (Carbonyl), 174.60 (Carbonyl), 174.39 (Carbonyl), 174.19 (Carbonyl), 174.13 (Carbonyl), 174.03 (Carbonyl), 100.38 ($C_1A_{major}$), 99.98 ($C_1A_{minor}$), 94.99 ($C_1B_{minor}$), 90.14 ($C_1B_{major}$), 79.24 ($_{minor?}$), 77.92 (CH-$Propionic_{minor}$), 77.40 (CH-$Propionic_{major}$), 76.13 ($C_3B_{major}$), 76.01 ($C_5$ $A_{major}$), 75.38 ($C_4B_{major}$), 75.08 ($_{minor?}$), 73.43 ($C_3A_{major}$), 70.92 ($C_5B_{major}$), 70.12 ($C_4A_{major}$), 61.39 ($C_6A_{major}$), 59.80 ($C_6B_{major}$), 59.62 ($_{minor?}$), 55.89 ($C_2A_{major}$), 54.86 (CH-$Lys_{major}$), 53.53 ($C_2B_{major}$), 52.87 (CH-D-$isogln_{major}$), 49.87 (CH-$ala_{major}$), 49.66 (CH-$ala_{minor}$), 39.12 ($CH_2$-$Lys\varepsilon_{major}$), 31.83 ($CH_2$-D-$isogln\gamma_{major}$), 30.90 ($CH_2$-$Lys\beta_{major}$), 27.09 ($CH_2$-D-$isolgn\beta_{major}$), 26.25 ($CH_2$-δ-$Lys_{major}$), 22.21(Acetate-$CH_{3minor}$), 22.07 (Acetate-$CH_{3major}$), 21.99 (Acetate-$CH_{3major}$), 21.96 ($CH_2$-$Lys\gamma_{major}$), 18.06 ($CH_3$-$Prop_{major}$), 17.86 ($CH_3$-$Prop_{minor}$), 16.64 ($CH_3$-$Ala_{minor}$), 16.55($CH_3$-$Ala_{major}$). HRMS m/z: $[M+H]^+$ calcd $C_{33}H_{58}O_{17}N_7^+$ 824.38837; Found 824.38872.

#### III. Bone Marrow Derived Macrophages culture and treatment

Bone marrow-derived macrophages (BMDMs) cells were generated by flushing bone marrow cells from femurs and tibia of mice, depleting red blood cells using ACK lysis buffer, and resuspending cell in complete DMEM media supplemented with 10% FBS, 0.5% penicillin/streptomycin mixture, and 20 ng/mL of M-CSF. Cells were maintained in culture at 37 °C, 5% $CO_2$ for 6 days before experimentation. BMDMs were stimulated with 20μM of designated compound dissolved in water. Water used as control treatment.

#### IV. Real-time quantitative-PCR and Western blotting

##### RNA isolation.

Total RNA from BMDMs was isolated using RNeasy Mini isolation kit (Qiagen) following the Qiagen Protocol (Part 1 and Part 2) using Ultrapure DNASE and RNASE Free water. The recommended-on column DNase digestion was performed during the isolation using RNase-Free DNase Set (Qiagen). Total RNA concentrations were determined using a nanodrop (Thermo Scientific).

**cDNA synthesis.**

cDNA synthesis was performed using a QuantiTect Reverse Transcription Kit (Qiagen). RNA was normalized prior to cDNA synthesis. Final concentrations of cDNA were determined, as well as 260/280 and 260/230 ratio, using a nanodrop (Thermo Scientific). Any cDNA with a 260/280 ratio below 1.8 was re-isolated.

**RT-PCR and Primers.**

Real-time PCR was performed in a 96-W plate (Multiplate™ Low-Profile 96-Well Unskirted PCR Plates #MLL9601 BIORAD) using a 7300 Applied Biosystems RT-PCR with SsoAdvanced Universal SYBR green Supermix (Bio-Rad).

PCR was performed with a reaction volume of 10 µL consisting of 0.3 µL each of F and R primer (10µM), 1 µL of cDNA (1:4 dilution), 5 µL of Sybr Green (Bio-Rad), and 3.4 µL of RNase/DNase Water. Each sample was repeated in duplicate for the PCR as a technical replicate. The thermal cycle was programmed for stage 1: 3 minutes at 95˚C; stage 2: followed by 40 cycles of 15 seconds at 95˚C and 45 seconds at 60˚C; stage 3: 10 seconds at 95˚C; stage 4: 10 seconds at 55˚C and 10 seconds at 95˚C.

Relative gene expression was determined utilizing the $2^{(-\Delta\Delta Ct)}$ method. Each compound treatment measurement in cells was normalized to the expression of the housekeeping gene 18s. Data was analyzed with GraphPad Prism 8 using an unpaired parametric t-test with equal SD two-tailed: * $P \leq 0.05$, **$P \leq 0.01$, ***$P \leq 0.001$, and ****$P \leq 0.0001$.

All primers were obtained from Eurofins Genomics LLC. A stock concentration of 100µM was made for each primer in RNase and DNase free water and stored at -80˚C. From the 100µM stocks, 10µM aliquots were made and stored at -20˚C for short term use.

***Cox2*_For:** CGAGTCGTTCTGCCAATAGAA
***Cox2*_Rev:** CCTGGTCGGTTTGATGTTACT
***Il-1β*_For:** GCAACTGTTCCTGAACTCAACT
***Il-1β*__Rev:** ATCTTTTGGGGTCCGTCAACT
***Oasl1*_1_For:** GTGCTCAAGGTACTCAAGGTAG
***Oasl1*_1_Rev:** CTGTGGAAACAGCTCAGGAA
***m18s*_For:** GTAACCCGTTGAACCCCATT
***m18s*_Rev:** CCATCCAATCGGTAGTAGCG
***tnfα*_For:** CAGGCGGTGCCTATGTCTC
***tnfα*_Rev:** CGATCACCCCGAAGTTCAGTAG
***Cxcl10*_For:** CCAAGTGCTGCCGTCATTTC
***Cxcl10*_Rev:** GGCTCGCAGGGATGATTTCAA

**PG library and results for qRT-PCR initial screen.**

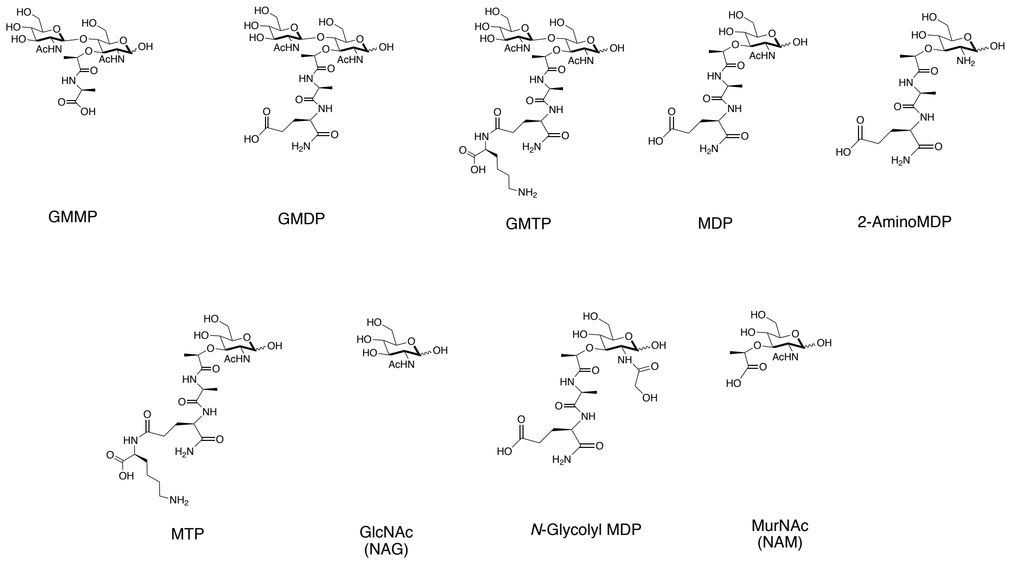


**SI-Figure 1:** Structures of PG synthetic fragments tested in qRTPCR assay. Compounds MDP (sigma-aldrich), GlcNAc (sigma-aldrich), *N*-glycolyl (ng-MDP) (invivogen), and MurNAc (sigma-aldrich) were purchased commercially. Compounds MTP and 2-Amino MDP were synthesized prior in our lab[1]. GMMP, GMDP, GMTP were synthesized and reported above.

### qRTPCR results for initial screen

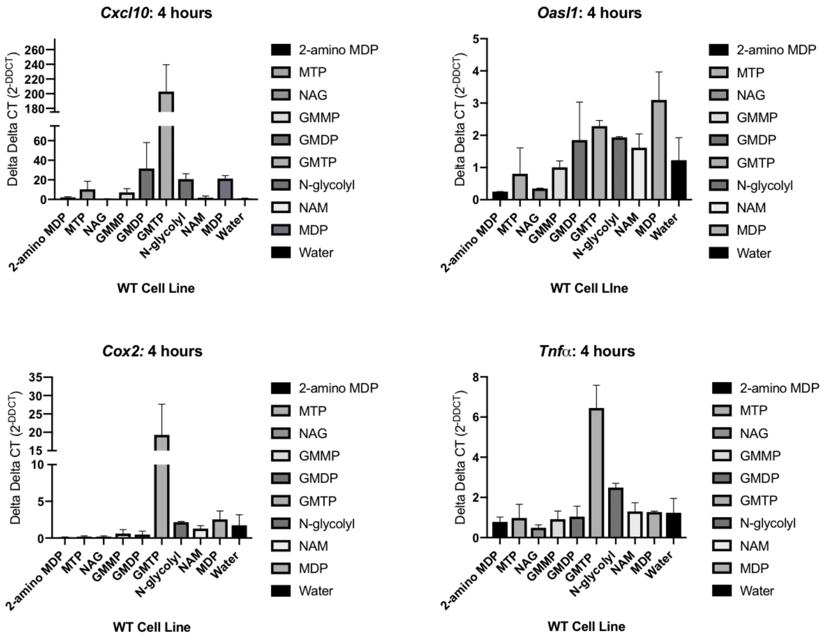


**SI-Figure 2**: qRTPCR cytokine screen of BMDMs treated with all PG fragments at 20 μM for 4 hours. Each compound was tested in biological duplicate with two technical replicates. Data was analyzed with GraphPad Prism 8. Error bars are reported as +/-SEM.

#### qRT-PCR GMTP vs. MDP 4-hour repeat.

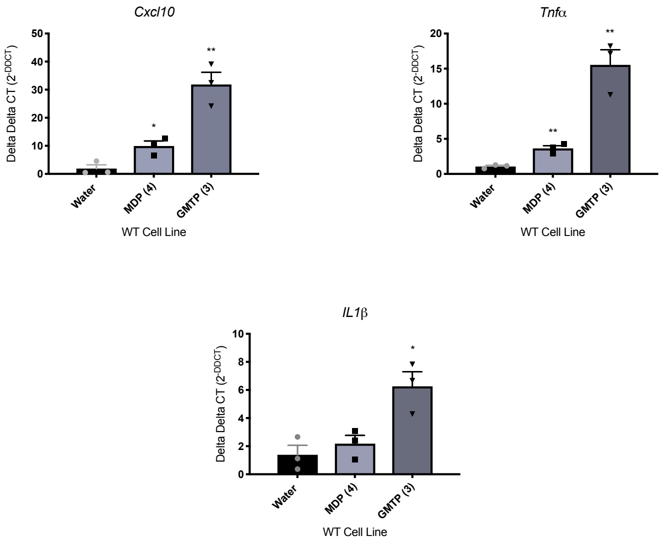


**SI-Figure 3:** Repeat in biological triplicate of BMDMs treated with 20μM of water, GMTP or MDP for 4 hours. Each compound/cytokine was tested via qRTPCR in two technical replicates. Data was analyzed with GraphPad Prism 8 using an unpaired parametric t-test with equal SD two-tailed: * P≤0.05, **P≤0.01, ***P≤0.001, and ****P≤0.0001. Error bars are reported as SEM.

**qRT-PCR Numerical ΔΔCT Data for Main Text Figure 3.**

**SI- Table 1: Individual replicate ΔΔCT for main Figure 3.**

| | Water | | | MDP (4) | | | N-glycolyl (ng-MDP) (5) | | | GMTP (3) | | |
|---|---|---|---|---|---|---|---|---|---|---|---|---|
| *Cxcl10* | 1.00000 | 1.06437 | 1.580083 | 13.45434 | 13.08643 | 6.916298 | 16.33619 | 18.89588 | 10.9283 | 20.9663 | 28.44297 | 27.66519 |
| *Tnfa* | 1.00000 | 1.166036 | 1.149521 | 2.892073 | 3.596983 | 3.304033 | 7.298441 | 5.776048 | 5.43299 | 15.06217 | 13.86763 | 13.62145 |
| *IL1b* | 1.00000 | 1.027679 | 1.181854 | 1.013861 | 1.087266 | 0.924596 | 4.224797 | 2.655366 | 2.70344 | 5.110092 | 3.927083 | 4.472197 |
| *Cox2* | 1.00000 | 1.043447 | 0.9427442 | 1.126976 | 1.157696 | 0.972567 | 3.788574 | 2.910659 | 2.67579 | 4.787842 | 4.500955 | 4.063975 |

#### V. <u>RNA sequencing</u>

**RNAseq protocols.**

BMDMs were generated from three WT mice and stimulated with 20 µM of different PG fragments in biological triplicate. After 18h of treatment total RNA was isolated using RNeasy Micro kit (Qiagen). RNA libraries were synthesized using Perkin Elmer's NEXTflex rapid directional RNAseq library preparation kit according to manufacturer's standard protocol (https://perkinelmer-appliedgenomics.com/wp-content/uploads/marketing/NEXTFLEX/RNA/5138-07_Rapid_Directional_RNA-Seq_Kit.pdf). The libraries were constructed on a Perkin Elmer Sciclone G3 NGS liquid handler. 310 ng of total RNA was used as input for all preparations. 15 PCR cycles were used for library amplification. Targeted sequencing depth was 30 million paired-end reads per sample performed in 3 biological replicates. Sequence data was generated on an Illumina NextSeq 550 system at the University of Delaware sequencing core.

**Data Analysis.**
Blc2fastq2 Conversion software (Illumina) was used to generate de-multiplexed Fastq files. Expression values were normalized as Fragments per Kilobase Million reads after correction for gene length (FPKM) in Cuffdiff version 1.05 in the DNAnexus analysis pipeline and filtered for genes that exhibited a statistically significant difference ($P<0.01$) with an FDR threshold of 0.05 and a biologically relevant change log fold change >1. Samples were analyzed in the RNA-sequencing pipeline of Seqmonk for mRNAs for opposing strand specific and paired end libraries with merged transcriptome isoforms, correction for DNA contamination and log transformed resulting expression values in $log_2$ FPKM. RNA sequencing data will be available at: https://www.ncbi.nlm.nih.gov/geo/query/acc.cgi?acc=.

#### VI. Biochemical Assays

##### Luminex.

Stored culture media samples (BMDMs treated with 20 µM compound or water control for 18h, as described in Section IV) were slowly thawed to room temperature and Luminex assay was performed at the University of Maryland, Baltimore Cytokine Core per manufacturer's instructions (Millipore):

> A 96 well plate (Greiner) was wet with 200 µl of Assay Buffer and placed on a shaker for 10 minutes. The plate is then decanted and 25 µl of Assay Buffer or appropriate buffer was added to each well and 25 µl of standard/sample/control was added to the appropriate wells. All samples were run in biological triplicate and technical duplicate. Then 25 µl of a mixture containing designated cytokines (1:50 dilution) that were conjugated to beads was added. All plates contain a minimum high and low control in order to determine the validity of the plates. The plate was then placed on a shaker, at 4°C overnight.
>
> The plate was then placed on a magnetic washer, 200 µl of Wash Buffer was added to each well, the plate was set on a shaker at 500 rpm for 1 minute and repeated an additional two times. After the last decanting step 25µl of detection antibody was added and the plate was placed on a shaker for one hour at room temp. Then 25 µl of Phycoerythrin (1:25 dilution) was added to each well and the plate was placed back on the shaker for 30 minutes. The plate was then washed three times and 150 ul of Sheath Fluid was added to each well. The plate was then read using a Luminex MagPix reader. The data was then calculated using Luminex's exponent Software.

##### Western Blot Analysis.

The following antibodies were used in this study: Rabbit antibodies against IRF5 obtained from Abcam (ab21689). Phospho-p65-NFκB (Ser536, 93H1), p65-NFκB (D14E12), STAT1 (D1K9Y) Phospho-STAT1 (Tyr701, 58D6). Mouse anti β-actin (8H10D10) were purchased from Cell Signaling Technology. The antibody against phospho-Ser445 IRF5 used at 1/2000 dilution was generated by immunizing rabbits with a synthetic peptide (IRLQIpS445NPDLC; NeoBiolab, MA. USA) and described[4]. Whole-cell extracts were obtained by harvesting cells with lysis buffer (1% NP-40, 20 mM Tris-HCl (pH 7.4), 150 mM NaCl, 2 mM EDTA, 2 mM EGTA, 4 mM $Na_3VO_4$, and 40 mM NaF) containing protease and phosphatase inhibitors tablets (Roche). Western blotting was performed using standard protocols for SDS-PAGE and wet transfer onto PVDF membranes. Primary antibodies were diluted in blocking buffer (BSA 5%+1X TBST) and incubated overnight at 4 °C. Secondary anti mouse (NA931V, GE Healthcare) or rabbit (NA934V, GE Healthcare) HRP were used at 1/5000 and incubated for 1 at RT. The bands were visualized by enhanced chemiluminescence (Western Lightning Plus [PerkinElmer]) and exposured on film.

#### VII. Peptidoglycan Enzymatic Degradation and Mass Spectrometry Analysis

***Lactobacillus acidophilus* growth and enzymatic degradation.**

*L. acidophilus* (Carolina Biological) was grown in an anaerobic environment in 95 mLs of Lactobacilli MRS Broth (10.0 g protease peptone #3, 10.0 g beef extract, 5.0 g yeast extract, 20.0 g dextrose, 1.0 g tween20, 2.0 g ammonium citrate, 5.0 g sodium acetate, 0.1 g magnesium sulfate, 0.05 g manganese sulfate, and 2.0 g dipotassium phosphate in 1L of water) for seven days at 37˚C.

After seven days, the cells were pelleted in a 50 mL conical tube at 3000 g for 5 minutes. Cells were washed with 10 mL of PBS and pelleted at 3000 g for 5 min two times. The pellet was resuspended in 7.5 mL of lysis buffer (50 mM Tris pH 8.0, 25 mM NaCl, 2 mM EDTA). Lysozyme from Chicken Egg White (Sigma) at 0.4 mg/ml was added and placed in shaker at 37˚C. Additional lysozyme was added twice a day for a total of six times over three days.

Digested samples were centrifuged for 30 s, and the supernatant was filtered through Amicon Ultra 3K Filter Devices. The filter was washed with water, and the resulting supernatants were combined and lyophilized to yield a clear oil (~75 mg).

**LC-Mass Spectrometry Method.**

A LC/MS method previously established by our lab, Liang and DeMeester et al, was utilized for testing a highly concentrated solution of isolated PG[6]. The eluting peaks were subjected to high-resolution mass analysis on the Q-Exactive Orbitrap (Acquity UPLC BEH C18 column 2.1 × 50 mm (Waters) using a Dionex UHPLC coupled to a Q-Exactive Orbitrap (Thermo Fisher Scientific)). The LC method was a 0.5 ml $min^{-1}$ linear gradient starting from 0% A to 50% B in 4 min. Eluent A was 0.1% formic acid in water and Eluent B was 0.1% formic acid in acetonitrile. Thermo Xcalibur Qual Browser was used to process and analyze the data generated. All species showed the expected mass, correct m/z ratio within ±10p.p.m. and correct isotopic pattern. All masses observed were not observed in the four negative controls utilizing same LC/MS method: (1) MRS growth media (1:25 dilution in water for injection), (2) PBS supernatant of the first and second washes of cells pellet before lysis buffer was added (direct injection), and (3) lysis buffer (1:25 dilution in water for injection).

#### Results and HRMS Spectra of Identified fragments.

**GM-Dipeptide**

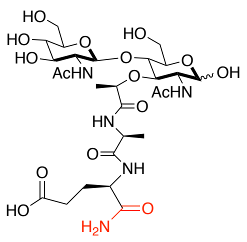


Chemical Formula: $C_{27}H_{45}N_5O_{16}$
Exact Mass: 695.28613

**GM-Tripeptide**

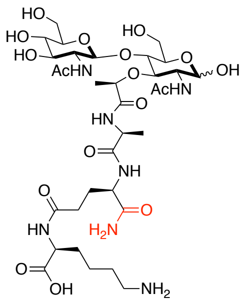


Chemical Formula: $C_{33}H_{57}N_7O_{17}$
Exact Mass: 823.38109

**GM-Tetrapeptide**

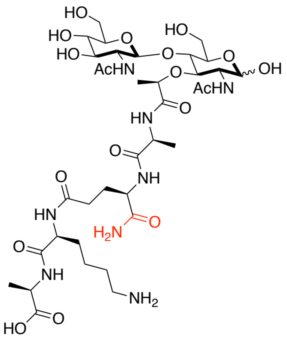


Chemical Formula: $C_{36}H_{62}N_8O_{18}$
Exact Mass: 894.41821

**GM-Pentapeptide**

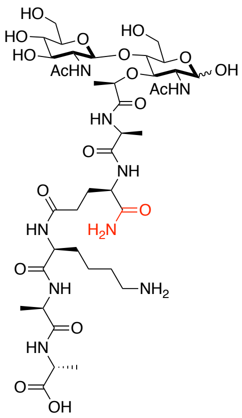


Chemical Formula: $C_{39}H_{67}N_9O_{19}$
Exact Mass: 965.45532

**SI-Table 2**: HRMS data obtained for each identified fragment along with its expected and observed mass. Repeated in three biological replicates (Observation/Replicate).

| Entry # | Fragment | Exact Mass | Expected [M+H] | Observed [M+H] | ppm | Retention Time (RT) | Ionization Intensity | Obs. /Rep. # |
|---|---|---|---|---|---|---|---|---|
| 1 | GM-Dipeptide | 695.28613 | 696.29341 | 696.29378 | 0.54155 | 2.53-2.58 | 1.59E+07 | 3/3 |
| **2** | **GM-Tripeptide** | **823.38109** | **824.38837** | **824.38770** | **-0.81394** | **1.44-1.48** | **2.65E+08** | **3/3** |
| 3 | GM-Tetrapeptide | 894.41821 | 895.42548 | 895.42461 | -0.97655 | 2.24-2.39 | 7.36E+07 | 3/ 3 |
| 4 | GM-Pentapeptide | 965.45532 | 966.46260 | 966.46363 | 1.06591 | 2.33-2.36 | 4.36E+06 | 3/3 |

#### ***Compound GM-Dipeptide (Entry #1)***

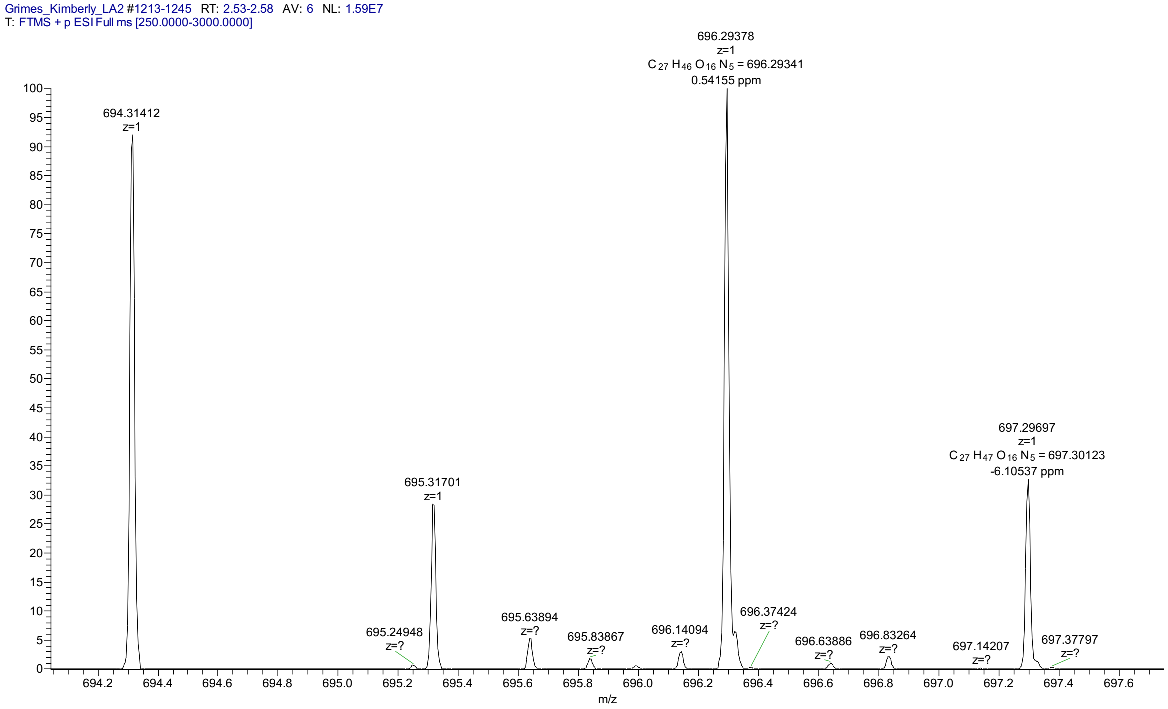


#### ***Compound GM-Tripeptide (Entry #2)***

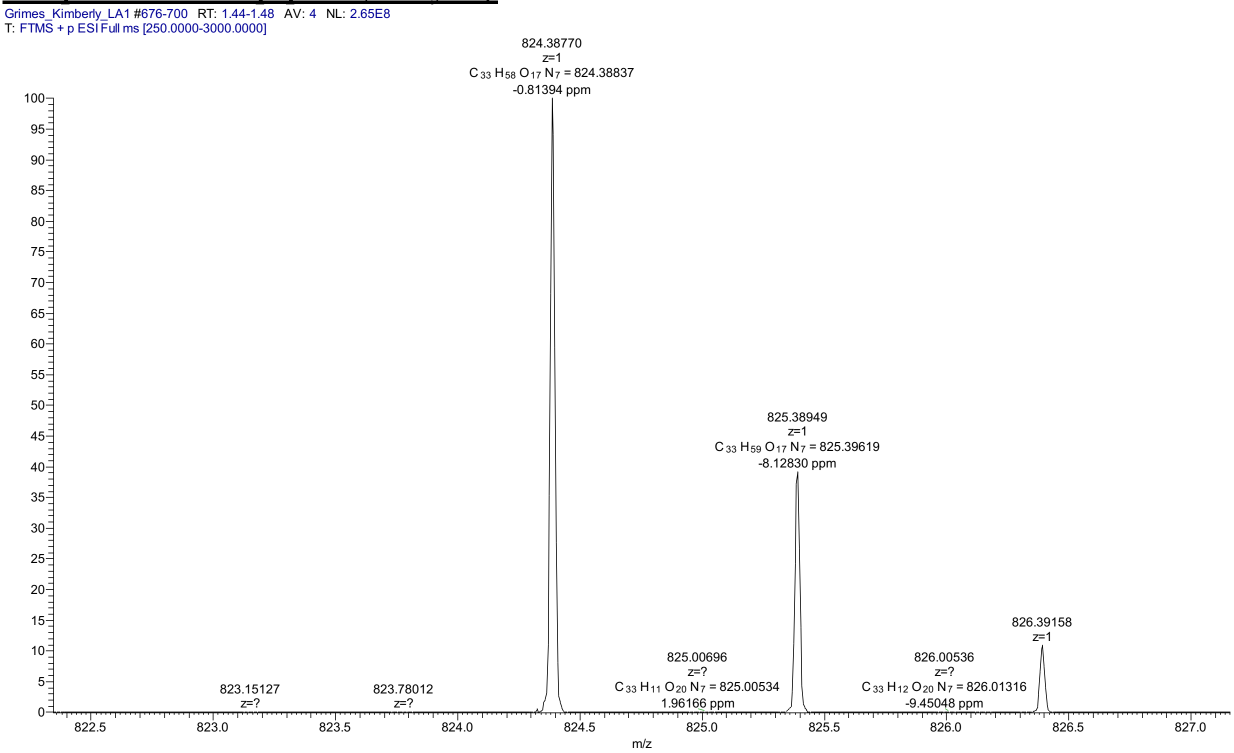

#### ***Compound GM-Tetrapeptide (Entry #3)***

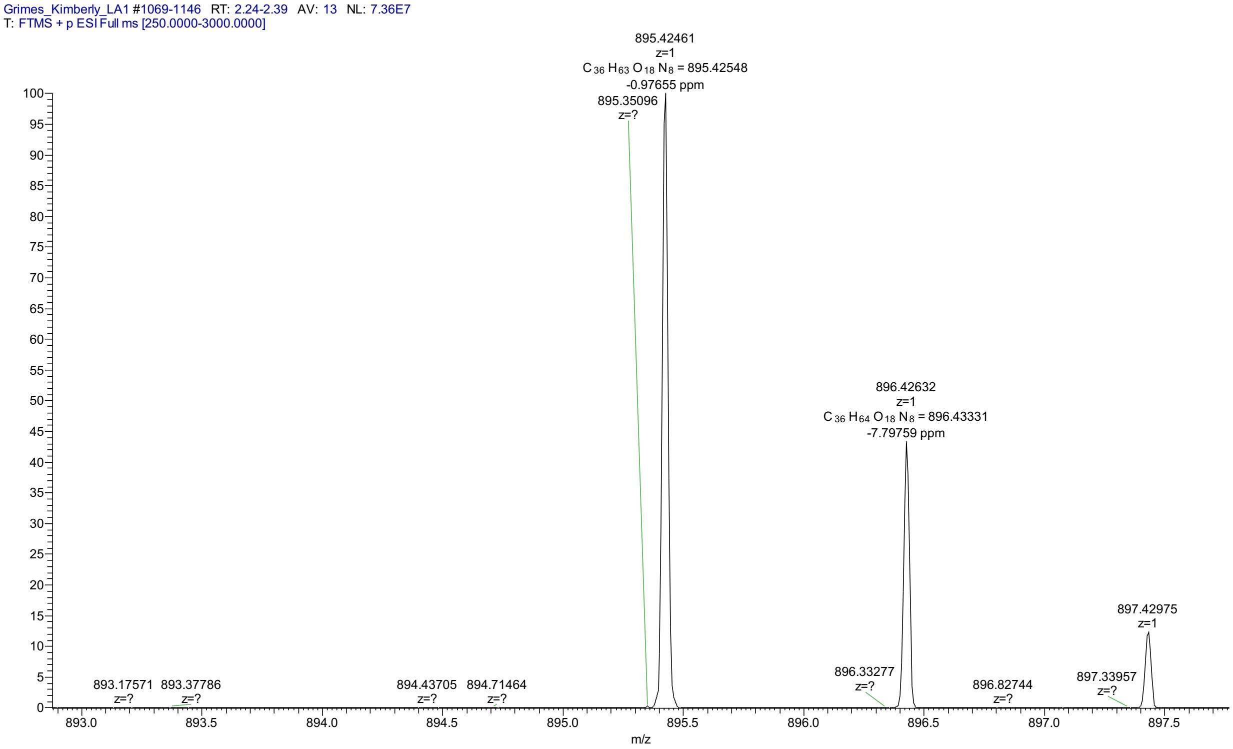


#### ***Compound GM-Pentapeptide (Entry #4)***

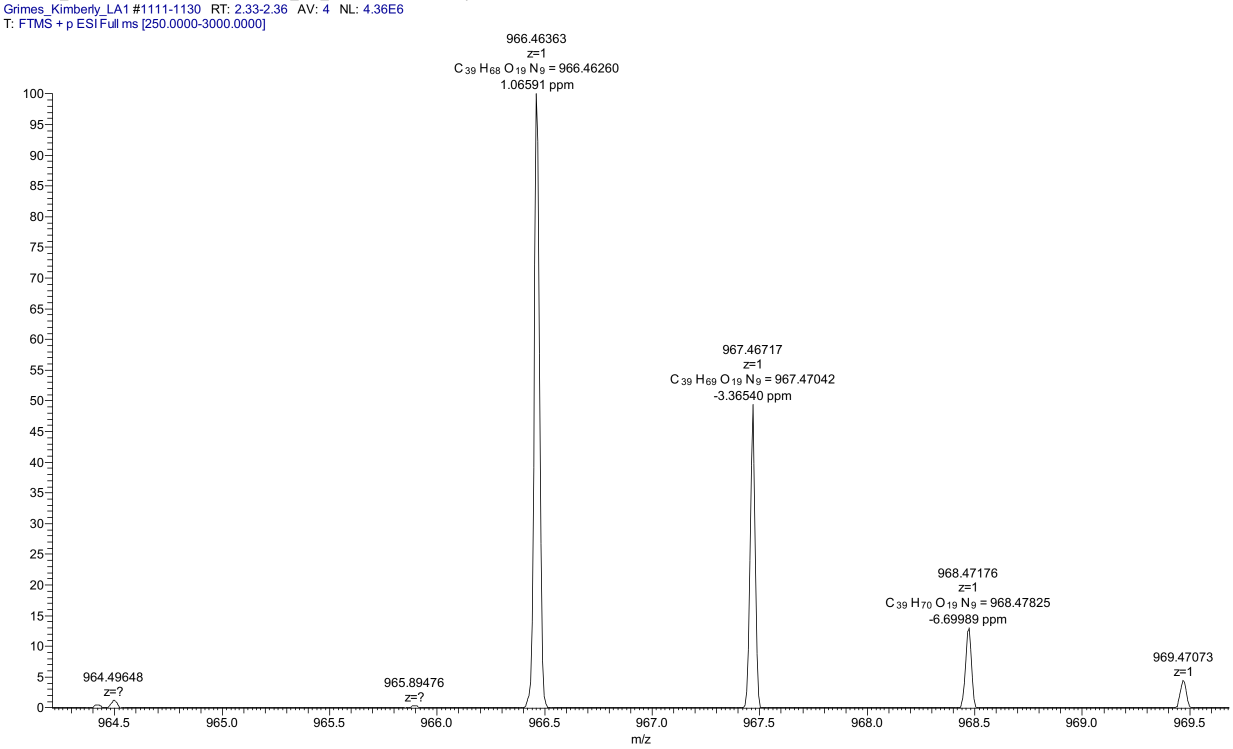

#### VIII. NMR Spectra of Synthetic Compounds

Compound **(1)** in $D_2O$

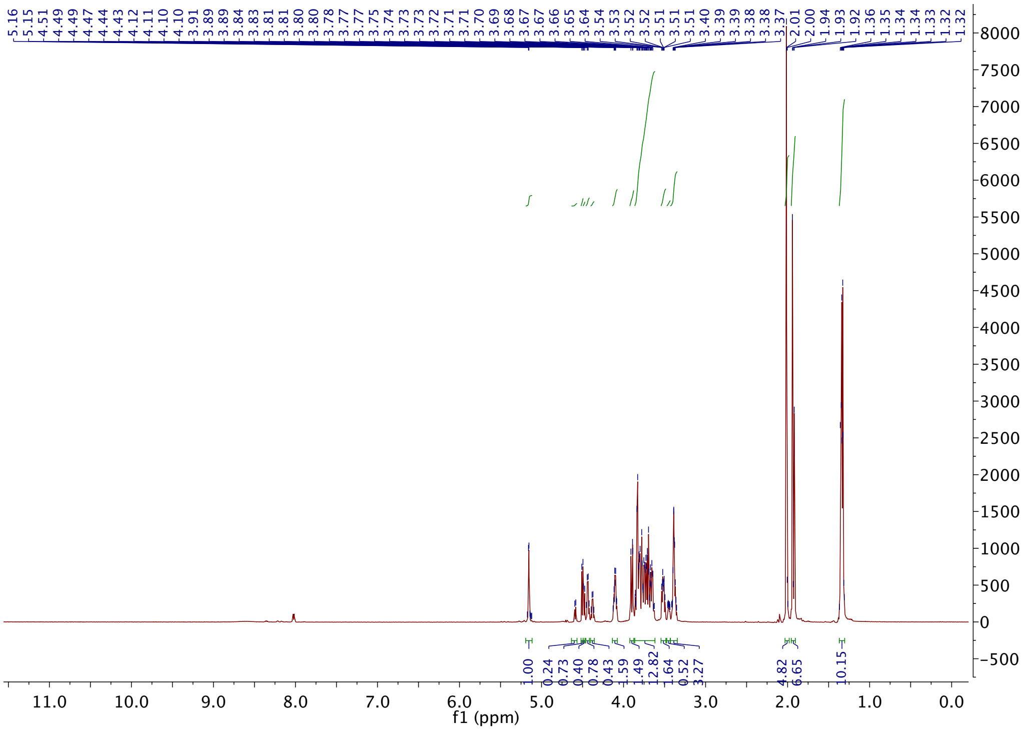


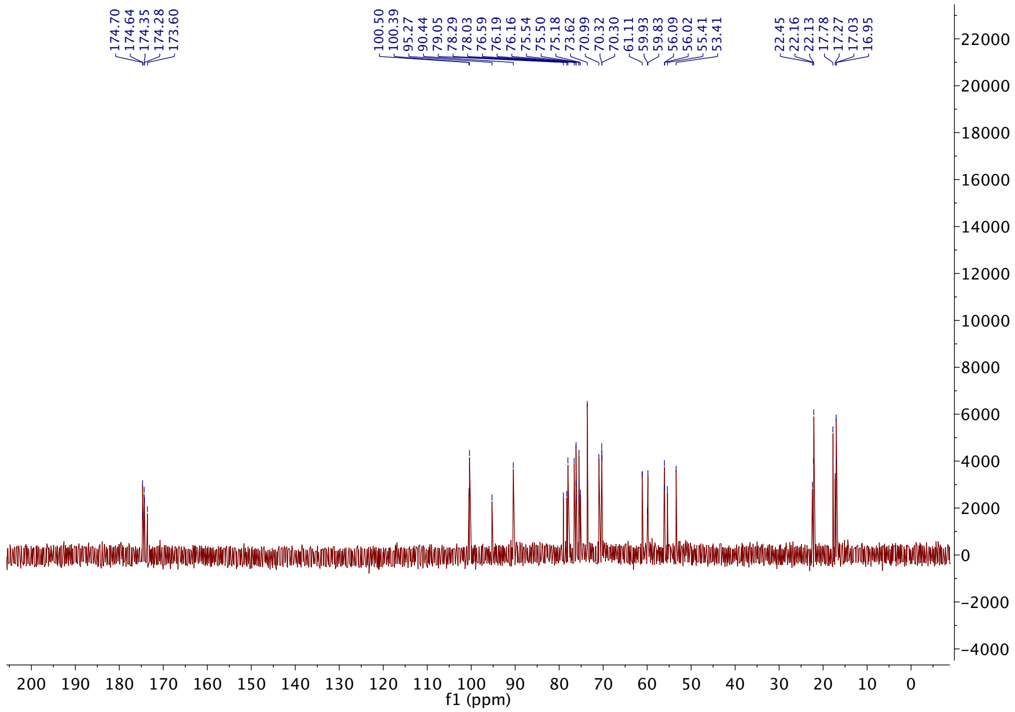

Compound (**11**) DMSO-*d6*

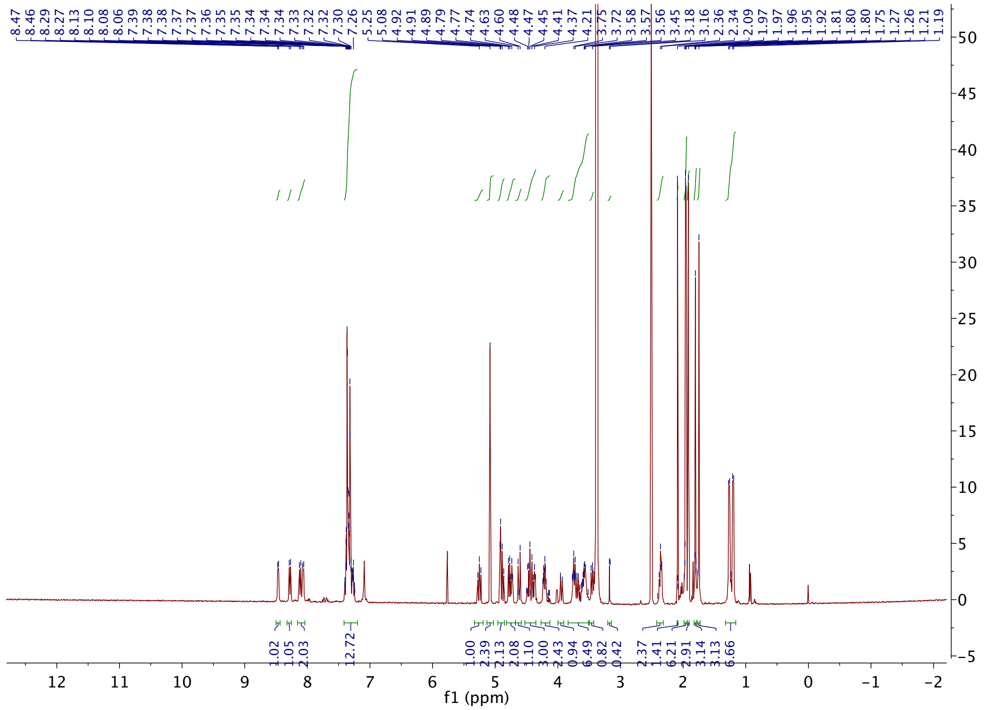

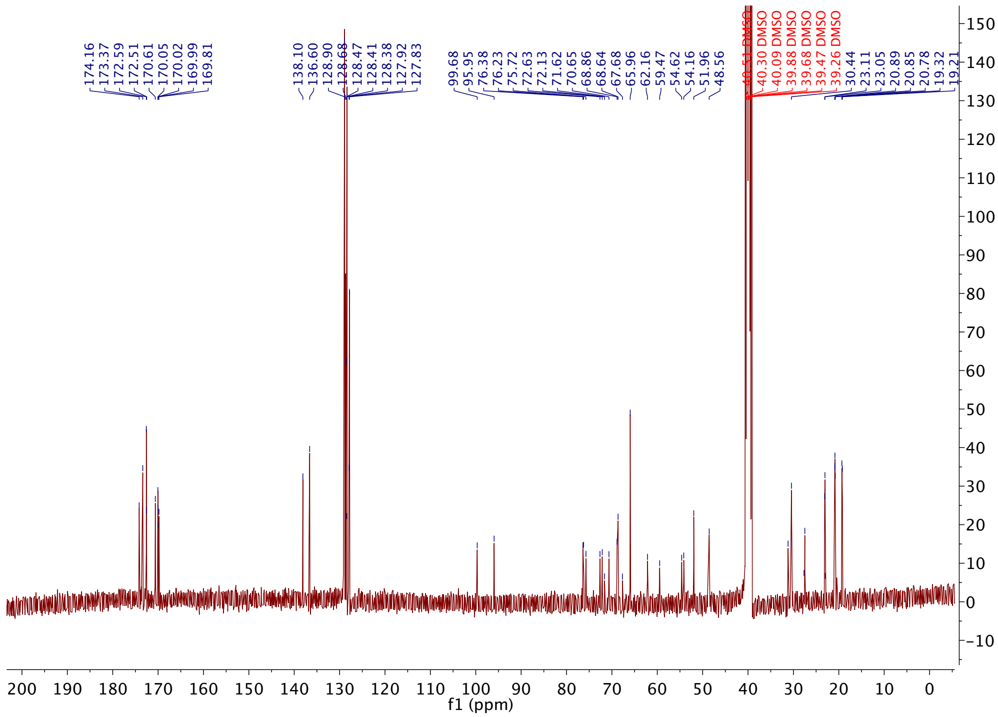

Compound (**2**) NMR in $D_2O$

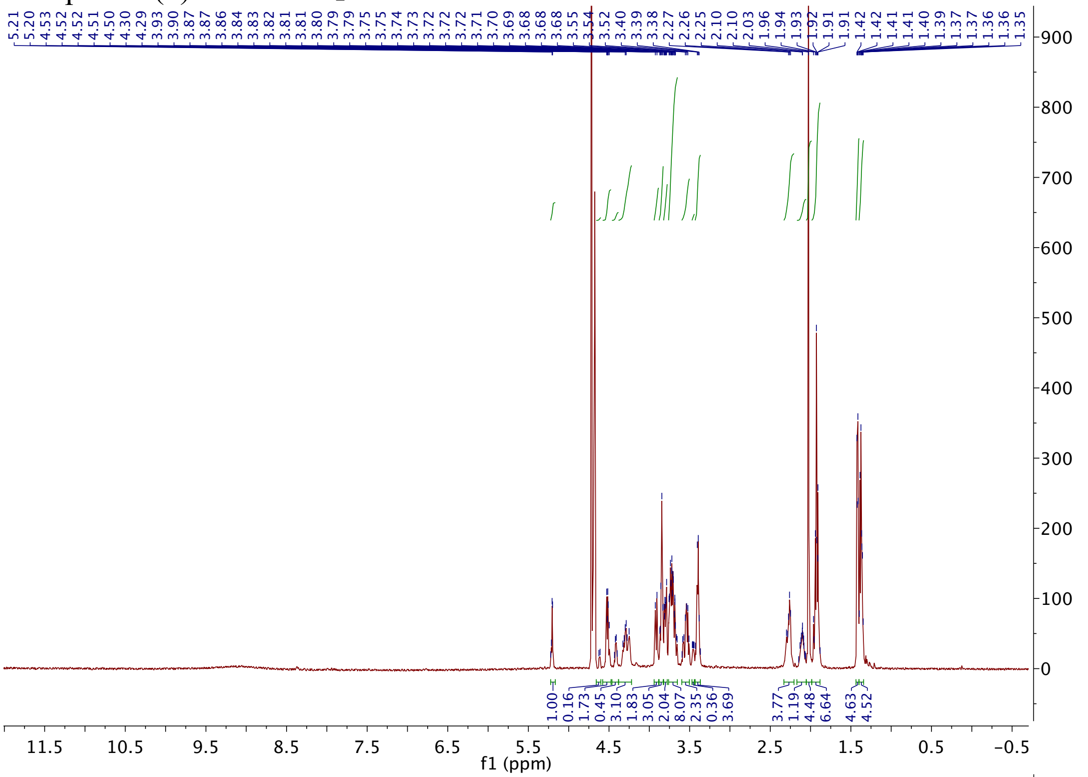

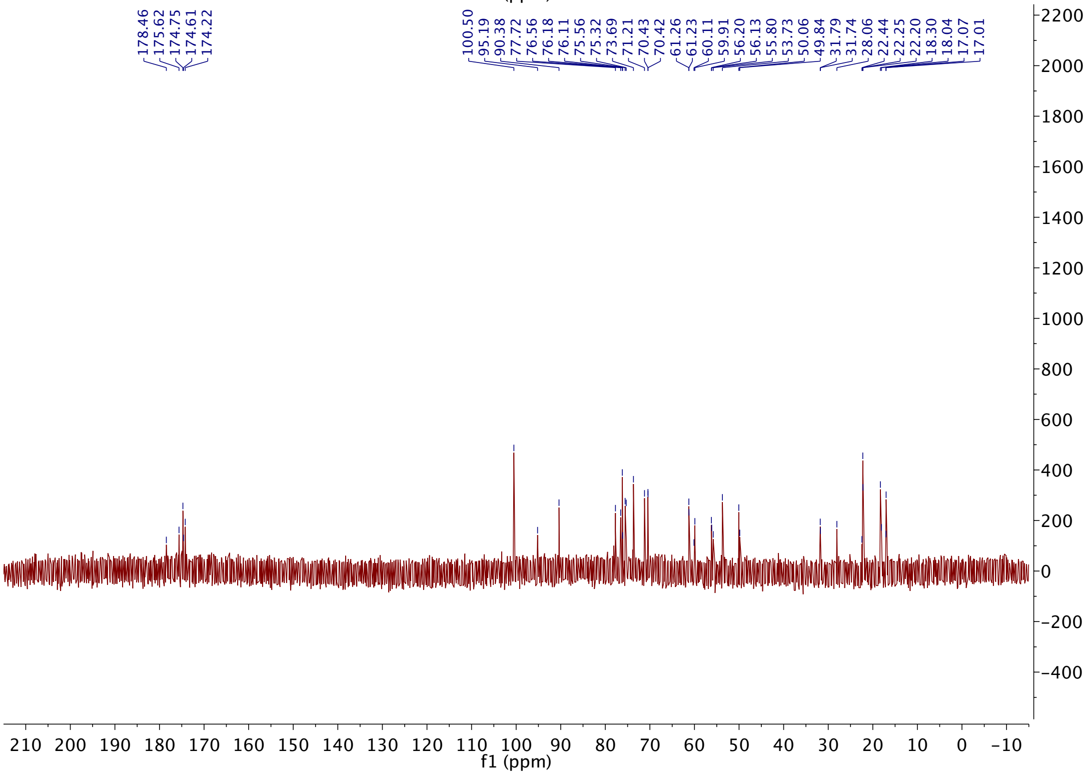

Compound (**12**) *DMSO-d6*

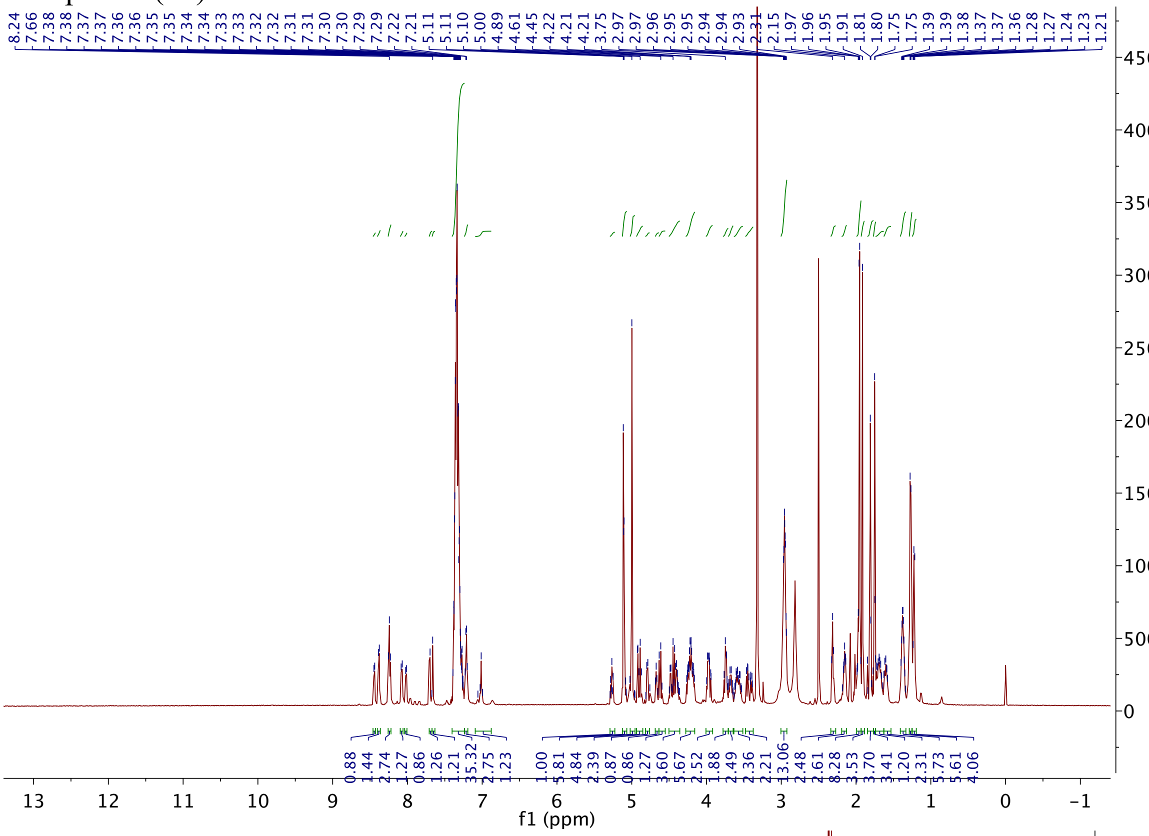

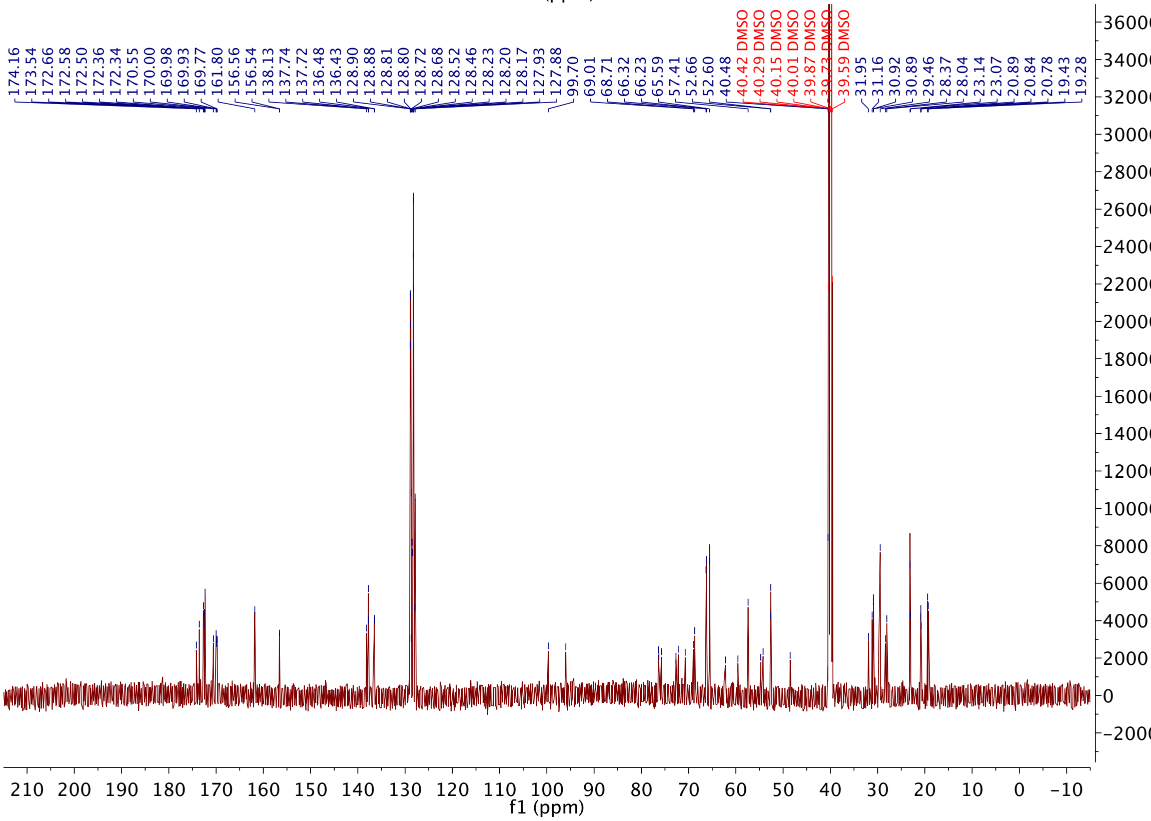

Compound (**3**) in $D_2O$

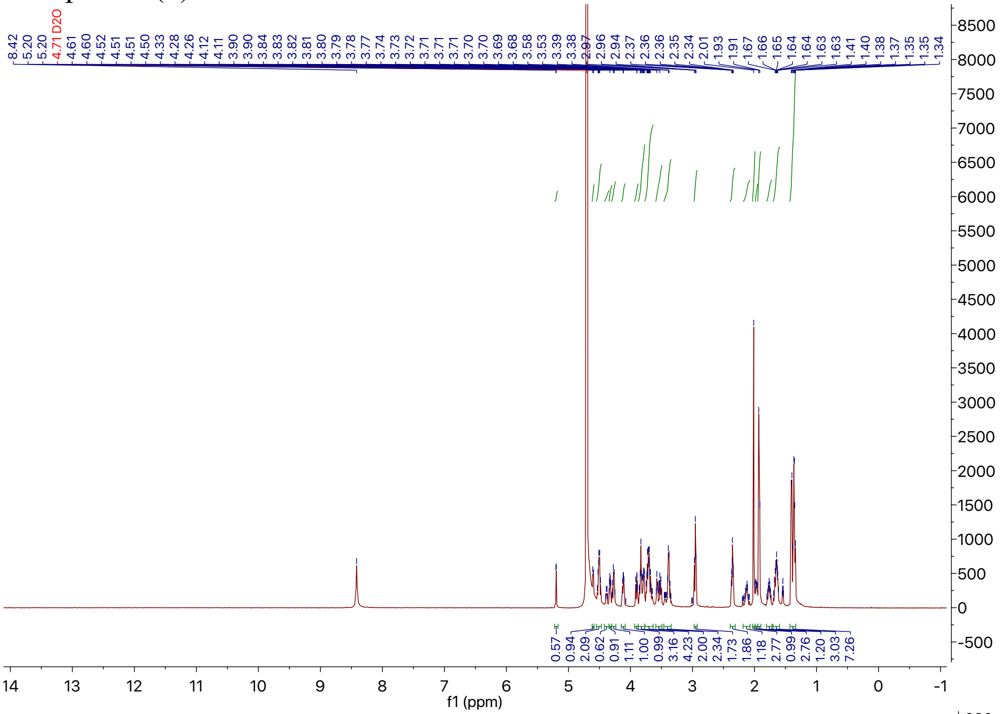

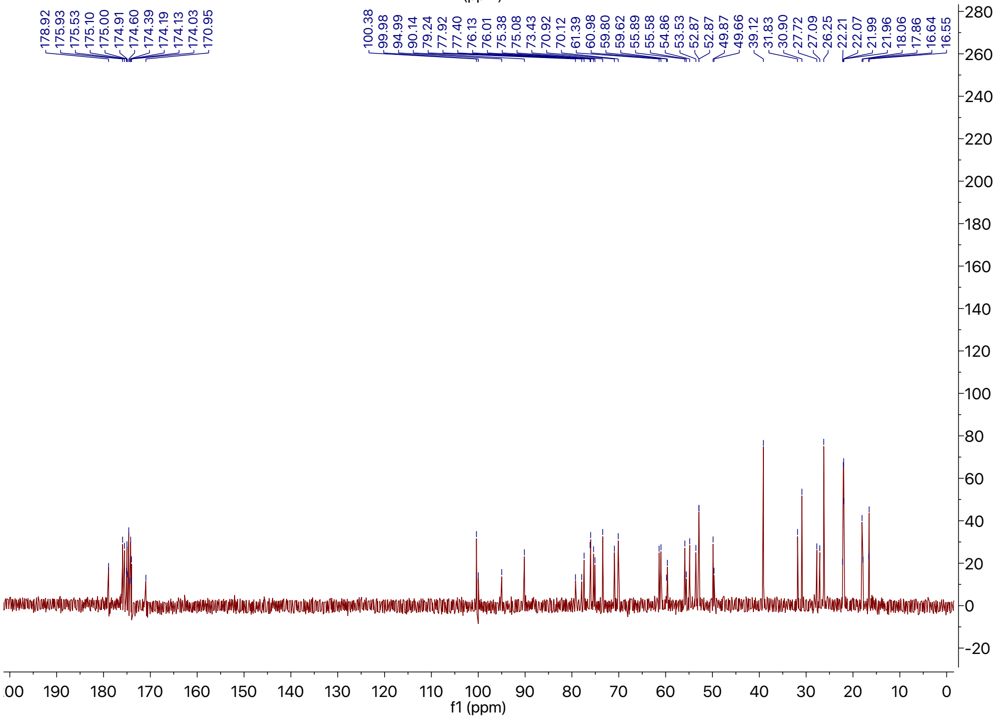

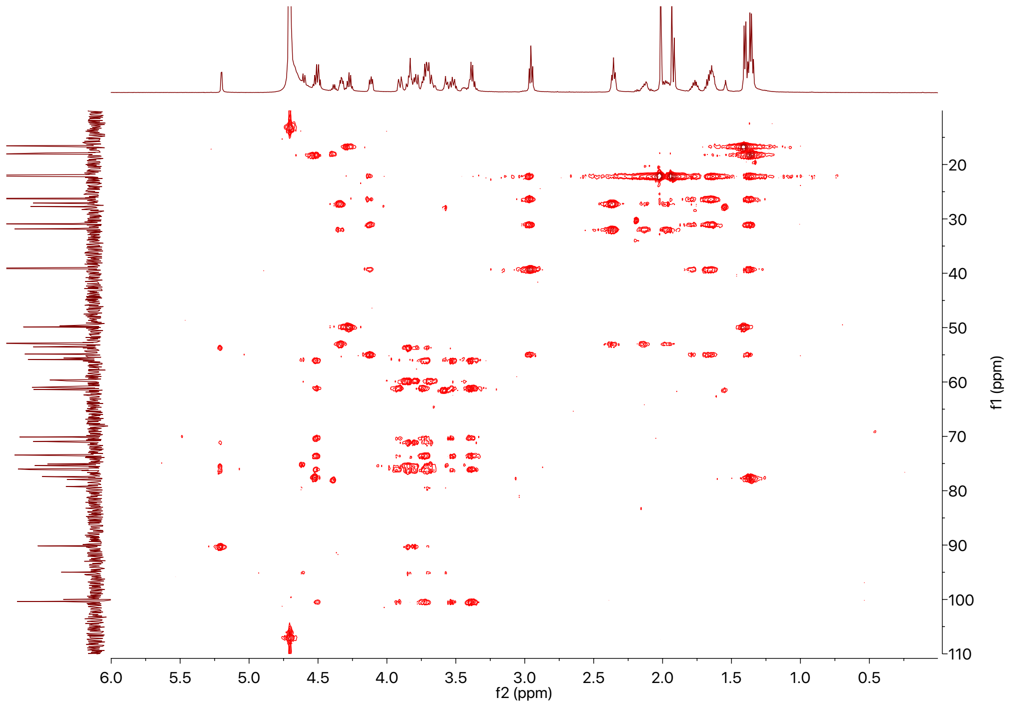
f1 (ppm)
f2 (ppm)

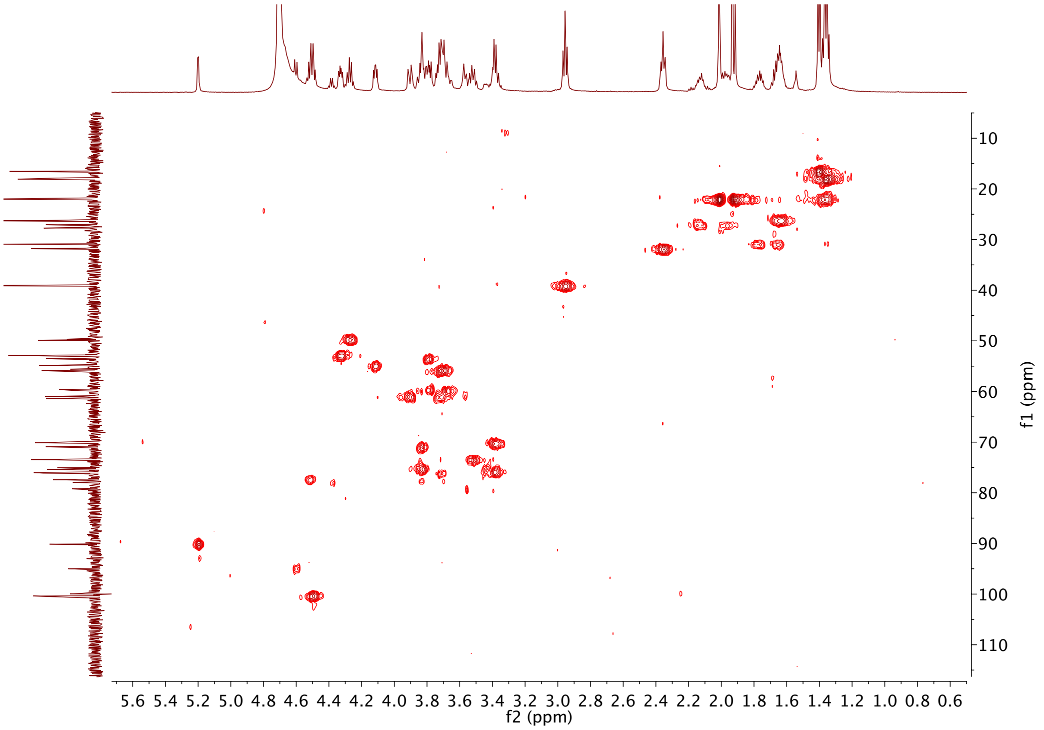
f1 (ppm)
f2 (ppm)

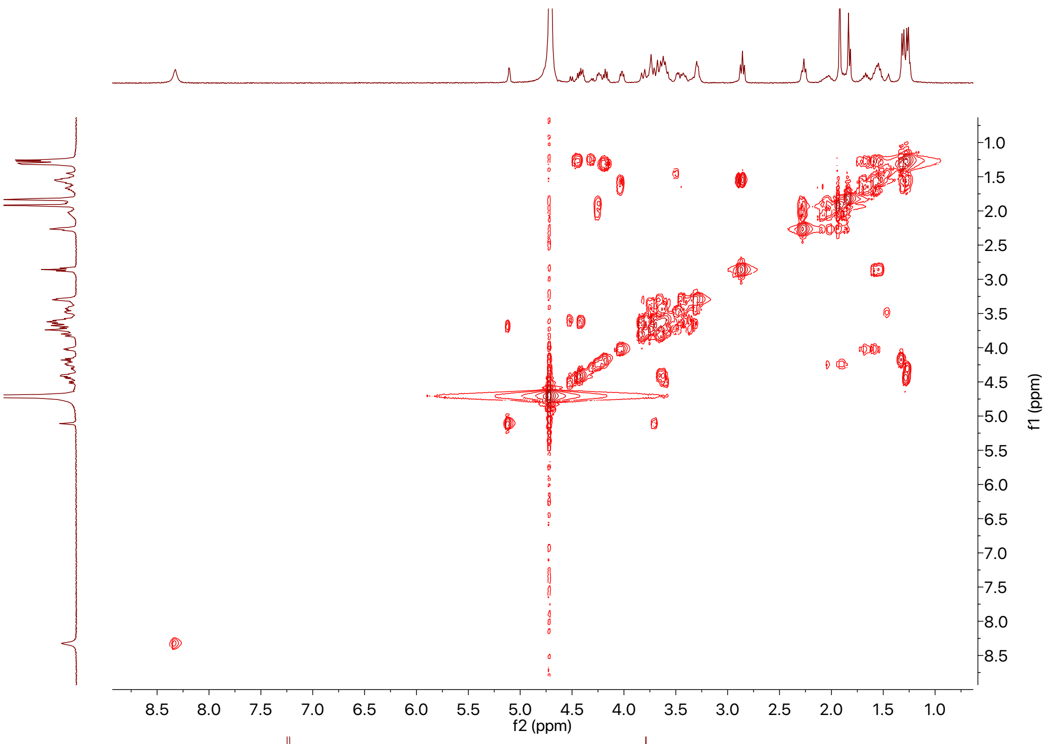
f2 (ppm)
f1 (ppm)

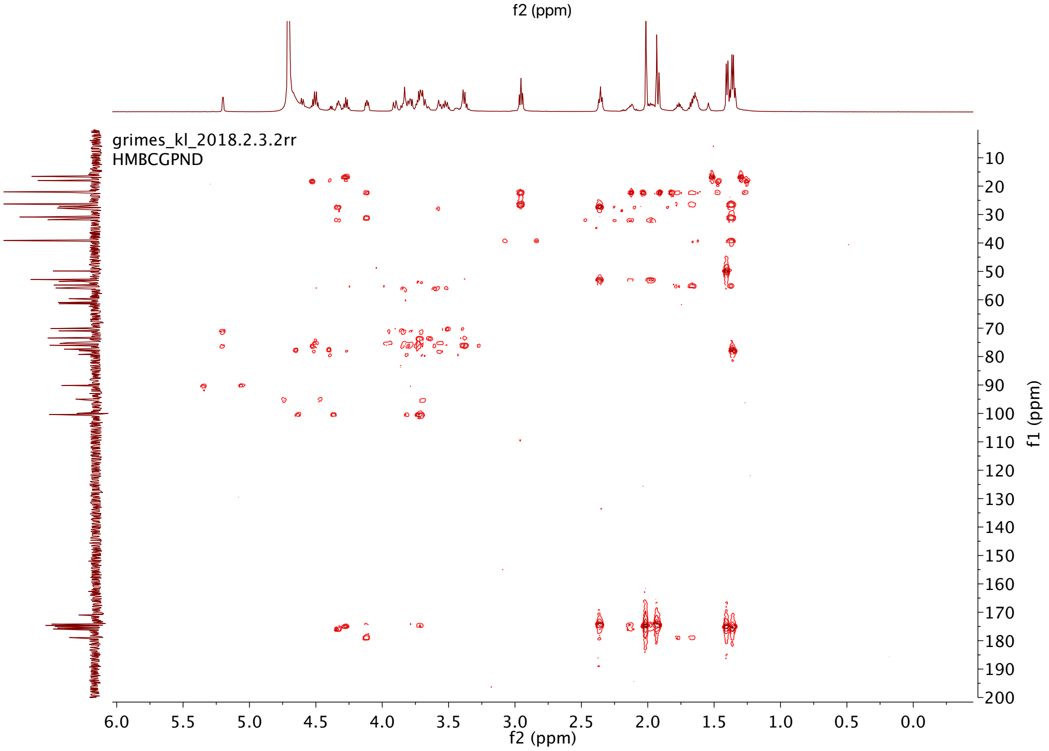
grimes_kl_2018.2.3.2rr
HMBCGPND
f2 (ppm)
f1 (ppm)

### IX. HRMS Spectra of Synthetic Compounds

#### Compound (**1**) HRMS Spectra

Grimes_Klare_KL-4-53B #40-82 RT: 0.21-0.36 AV: 4 NL: 6.26E7
T: FTMS + p ESI Full ms [100.00-1500.00]

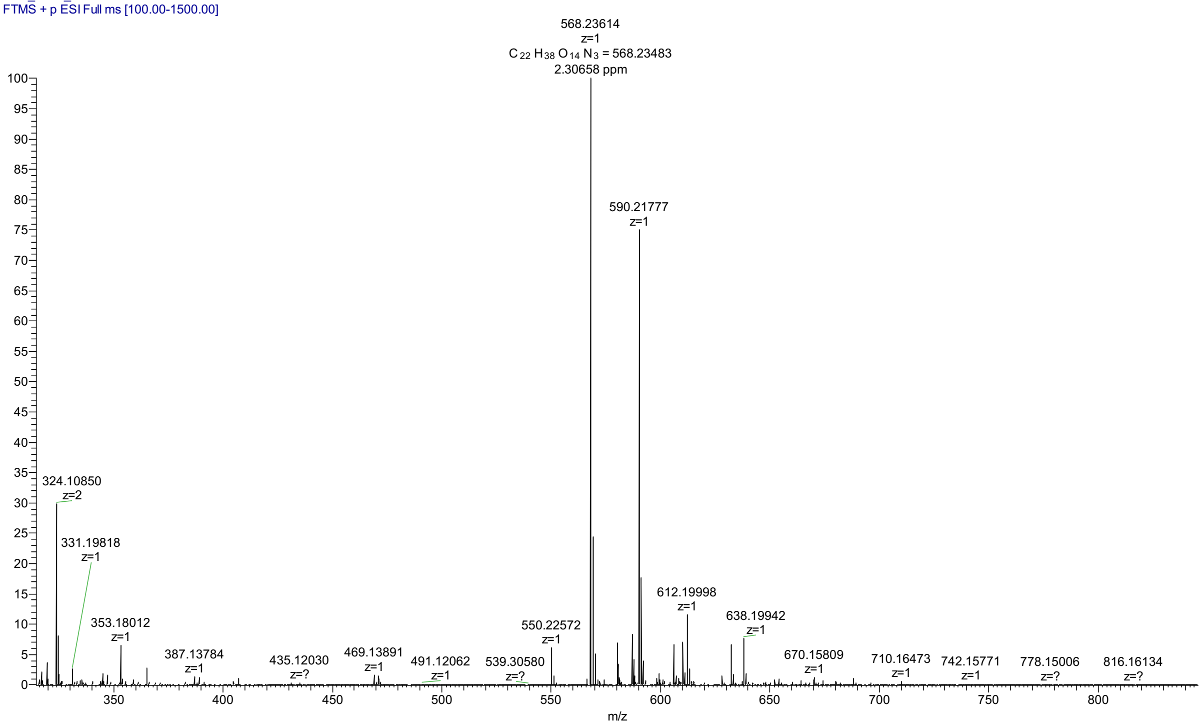


#### Compound (**11**) HRMS Spectra

Grimes_Klare_KL-4-46 #50-67 RT: 0.26-0.31 AV: 2 NL: 1.91E8
T: FTMS + p ESI Full ms [100.00-1500.00]

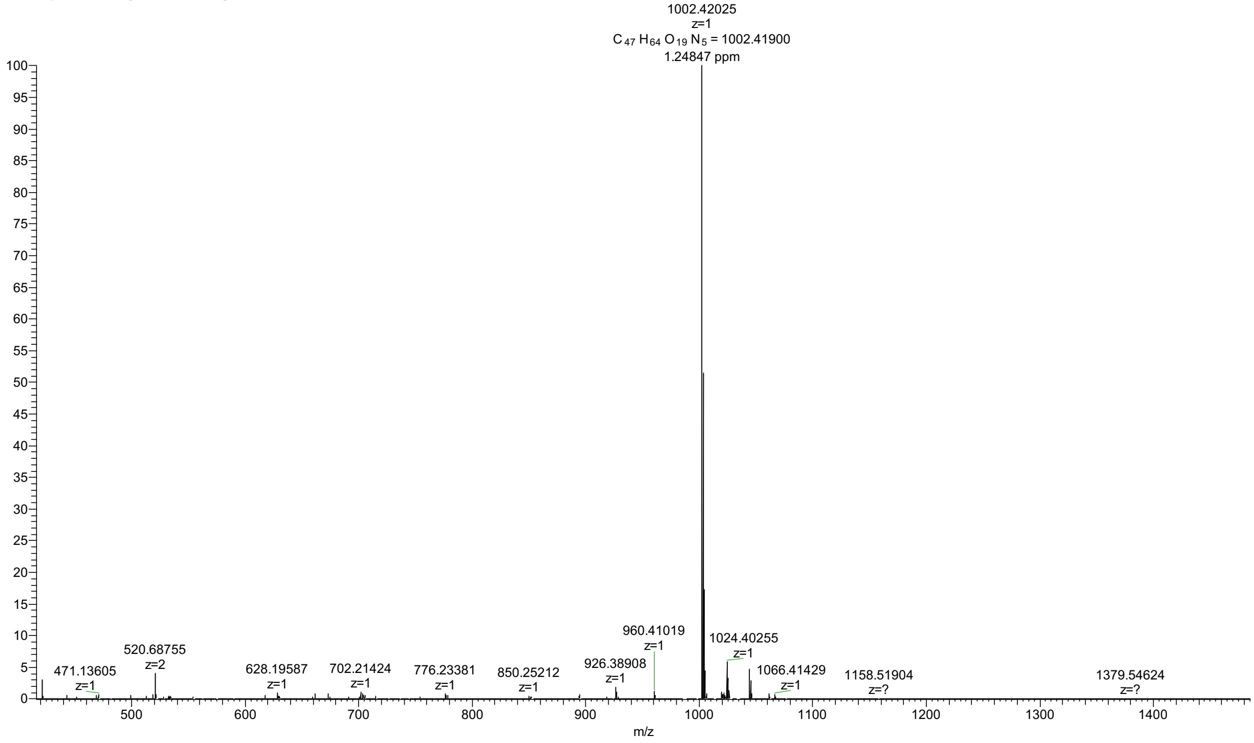

#### Compound (**2**) HRMS Spectra

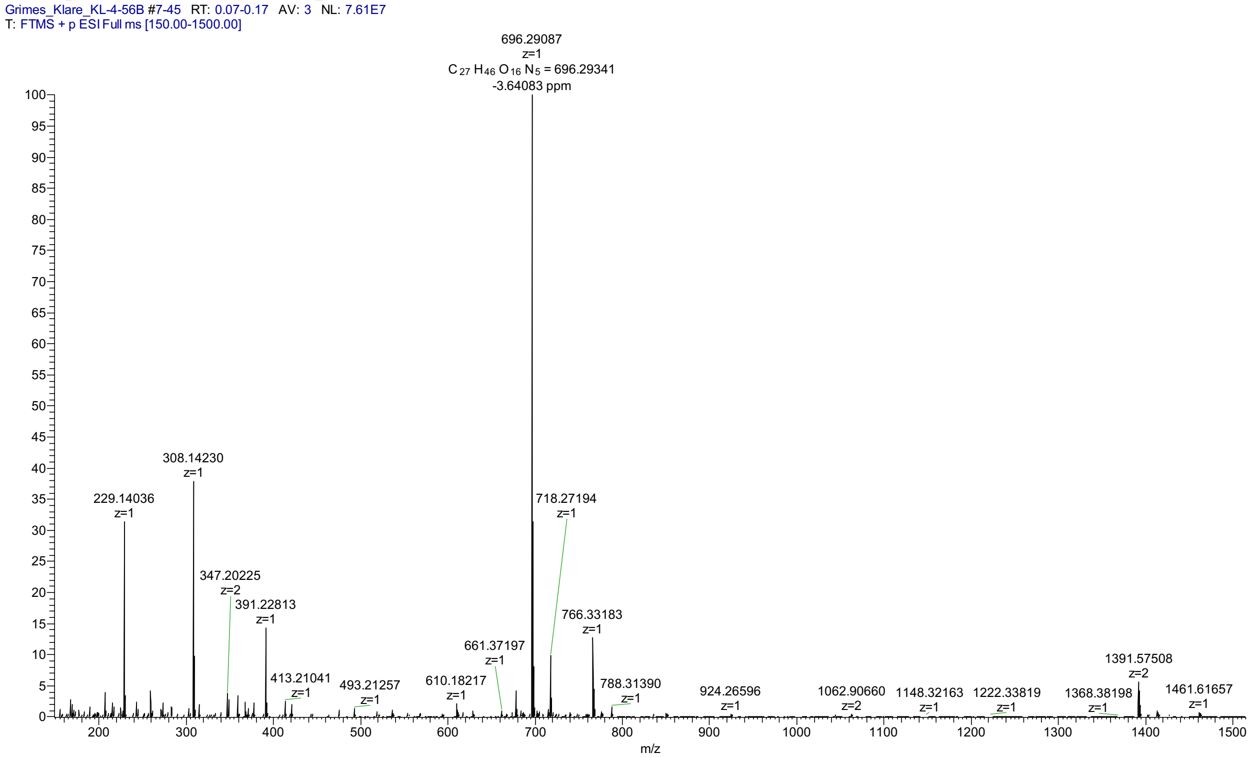


#### Compound **(12)** HRMS Spectra

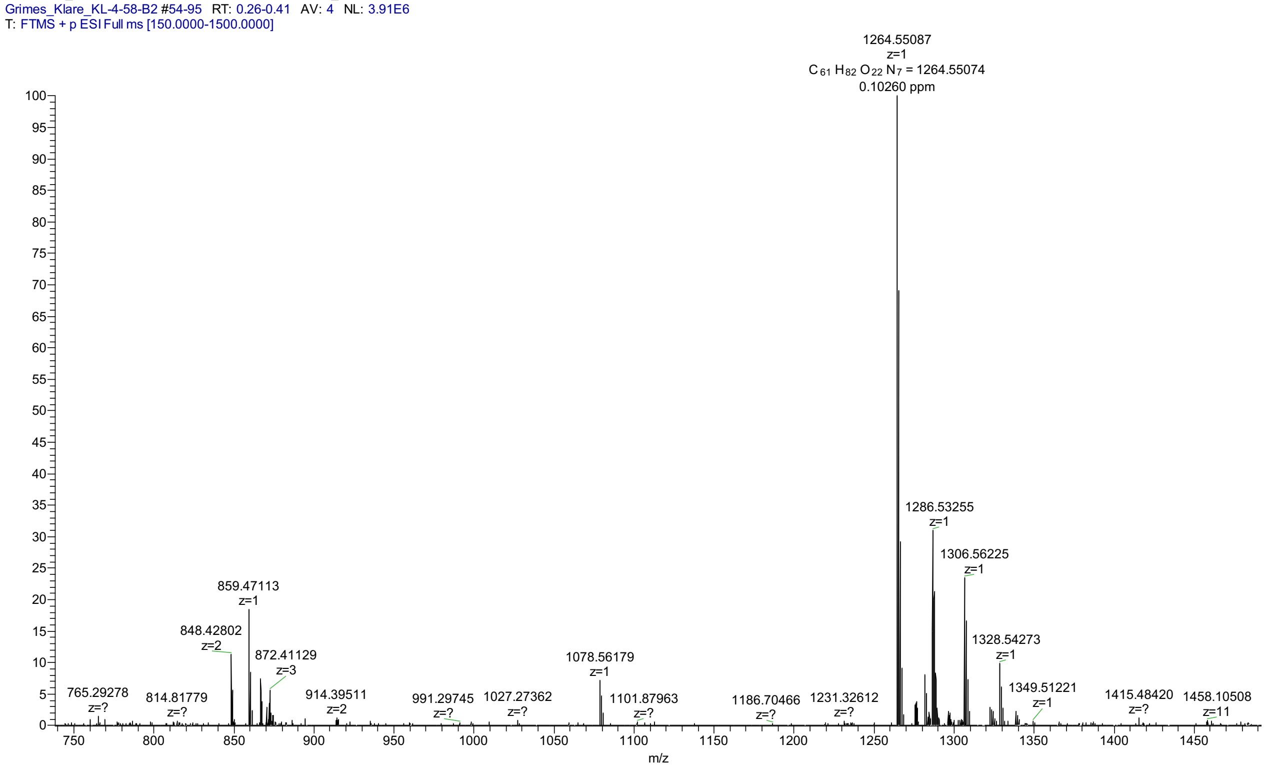

Compound (**3**) HRMS Spectra

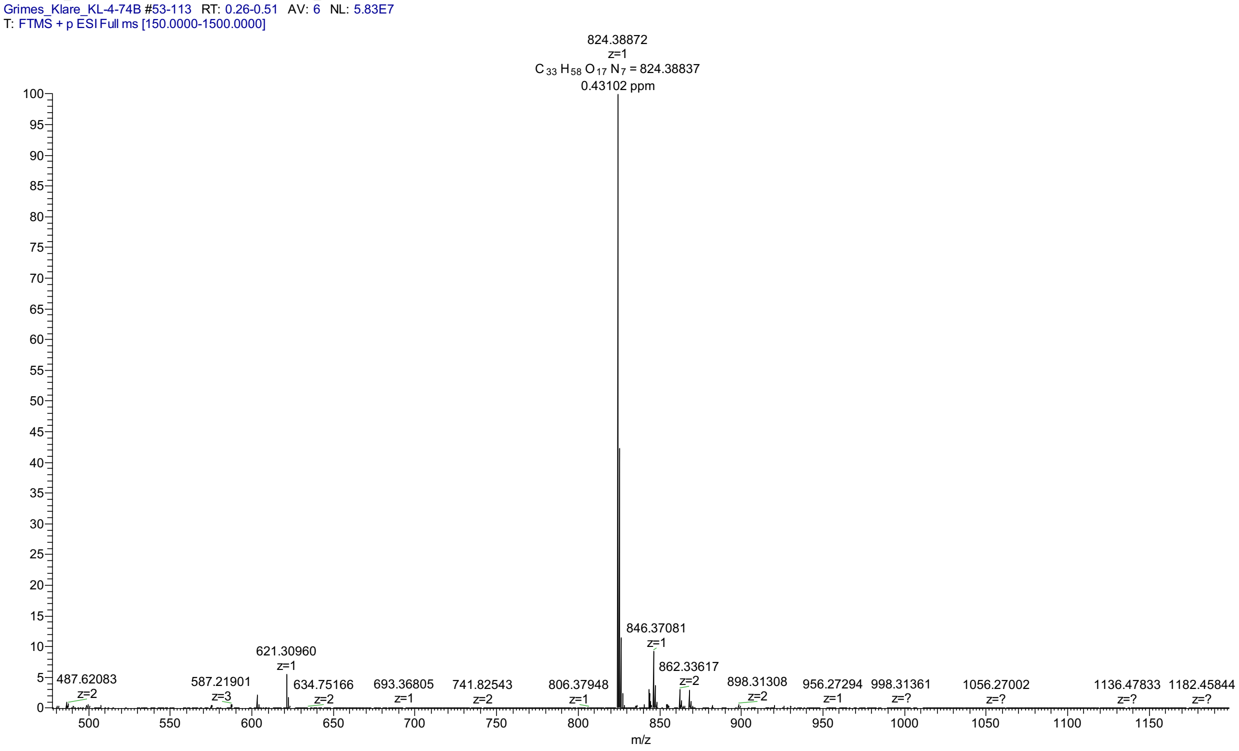


#### X. <u>References</u>
